# Prediction of protein subcellular localization in single cells

**DOI:** 10.1101/2024.07.25.605178

**Authors:** Xinyi Zhang, Yitong Tseo, Yunhao Bai, Fei Chen, Caroline Uhler

## Abstract

The subcellular localization of a protein is important for its function and interaction with other molecules, and its mislocalization is linked to numerous diseases. While atlas-scale efforts have been made to profile protein localization across various cell lines, existing datasets only contain limited pairs of proteins and cell lines which do not cover all human proteins. We present a method that uses both protein sequences and cellular landmark images to perform **P**redictions of **U**nseen **P**roteins’ **S**ubcellular localization (**PUPS**), which can generalize to both proteins and cell lines not used for model training. PUPS combines a protein language model and an image inpainting model to utilize both protein sequence and cellular images for protein localization prediction. The protein sequence input enables generalization to unseen proteins and the cellular image input enables cell type specific prediction that captures single-cell variability. PUPS’ ability to generalize to unseen proteins and cell lines enables us to assess the variability in protein localization across cell lines as well as across single cells within a cell line and to identify the biological processes associated with the proteins that have variable localization. Experimental validation shows that PUPS can be used to predict protein localization in newly performed experiments outside of the Human Protein Atlas used for training. Collectively, PUPS utilizes both protein sequences and cellular images to predict protein localization in unseen proteins and cell lines with the ability to capture single-cell variability.

## Introduction

The subcellular localization of a protein is important for its function and interaction with other molecules; its mislocalization is directly associated with various diseases^1–3^. In fact, a significant number of diseases with comorbidities could be better explained by protein subcellular localization than shared genes or protein-protein interactions^2^. Atlas-scale efforts have been made to spatially profile thousands of proteins in multiple cell lines through immunofluorescence microscopy^3^, live-cell imaging^4^, and mass spectrometry^5^. However, the number of possible combinations of proteins and cell lines far exceeds the number of combinations that have been measured even in the largest subcellular localization datasets^3,5^. For example, while the Human Protein Atlas (HPA) provides the subcellular localization of proteins encoded by 13,147 genes, this is only 65% of all known human protein-coding genes, and each protein is only measured in up to three cell lines with a total of 37 cell lines in the entire dataset^3^ (Figure 2a). Additionally, experimental staining of proteins is limited by the number of proteins that can be simultaneously measured in the same cell, with a typical multiplexing experiment allowing the staining of around 30 proteins in fixed samples^6^.

Furthermore, given the observed variability of protein localization not just across cell lines but also across single cells within a cell line^3^, the localization measured for a particular protein and cell line pair in an existing atlas might not be applicable to a newly performed experiment with different biological context that might change protein localization.

Recent advances in machine learning have enabled the study of protein properties using either protein sequences or cellular images. Models have been developed that infer different properties of a protein, such as its localization and structure, from the protein sequence^7–12^.

While the prediction of protein localization based on its sequence allows generalizing to unseen proteins, the localization prediction task cannot capture relative protein abundance among different cellular compartments, contextual differences in localization among single cells, or cell-type specific localization differences among cell lines. In addition, both supervised and unsupervised learning models have been developed to either automatically annotate or predict the cellular compartments that a particular protein localizes to using protein images^13–15^. While some of these models can reveal single-cell variability in protein localization^14,15^, these models for cellular compartment annotation require actual measurements of the protein images and cannot be used to predict protein localization of unseen proteins or in unmeasured cells. Given the large number of proteins and cell lines not measured in existing atlases and the known cell type variability in protein localization, new computational models are needed to predict protein localization at single-cell level for proteins and cell lines that have not been used for training the computational model.

We present a method that uses both protein sequences and cellular landmark images to perform **P**redictions of **U**nseen **P**roteins’ **S**ubcellular localization (**PUPS**), which can generalize to proteins and cell lines not used for model training. PUPS combines a protein language model and an image inpainting model to utilize both protein sequence and cellular images of landmark stains for predicting protein localization. These two ingredients allow our model to overcome the above mentioned limitations of prior approaches: The protein sequence input allows generalization to unseen proteins and the landmark stains allow cell type specific predictions that capture single-cell variability and can generalize to unseen cell lines (Figure 1b). We demonstrate that our model has low prediction error, even when both the protein and the cell line were not used in training. The ability of PUPS to generalize to unseen proteins and cell lines enables us to assess the variability in protein localization across cell lines as well as across single cells within a cell line, which has been difficult given that existing atlases do not fully cover all protein and cell line pairs (Figure 1a, 2a). We identified that the proteins with the most variable ratios between nucleus and cytosol across cell lines are associated with transcription, cell differentiation, and chromatin regulation, whereas high single-cell variability in protein localization within a cell line is mainly associated with cell division, transcription, double-strand break repair, and apoptosis. We performed imaging experiments to validate that PUPS can be used for protein localization prediction in experiments outside the HPA used to train our model, and we showed that the protein localization variability based on the predicted images is accurate. Finally, we provide intuition for PUPS’s generalization ability and show that PUPS is able to extract known protein sequence features relevant to localization as well as learn meaningful representations of proteins and cell lines.

**Figure 1.**
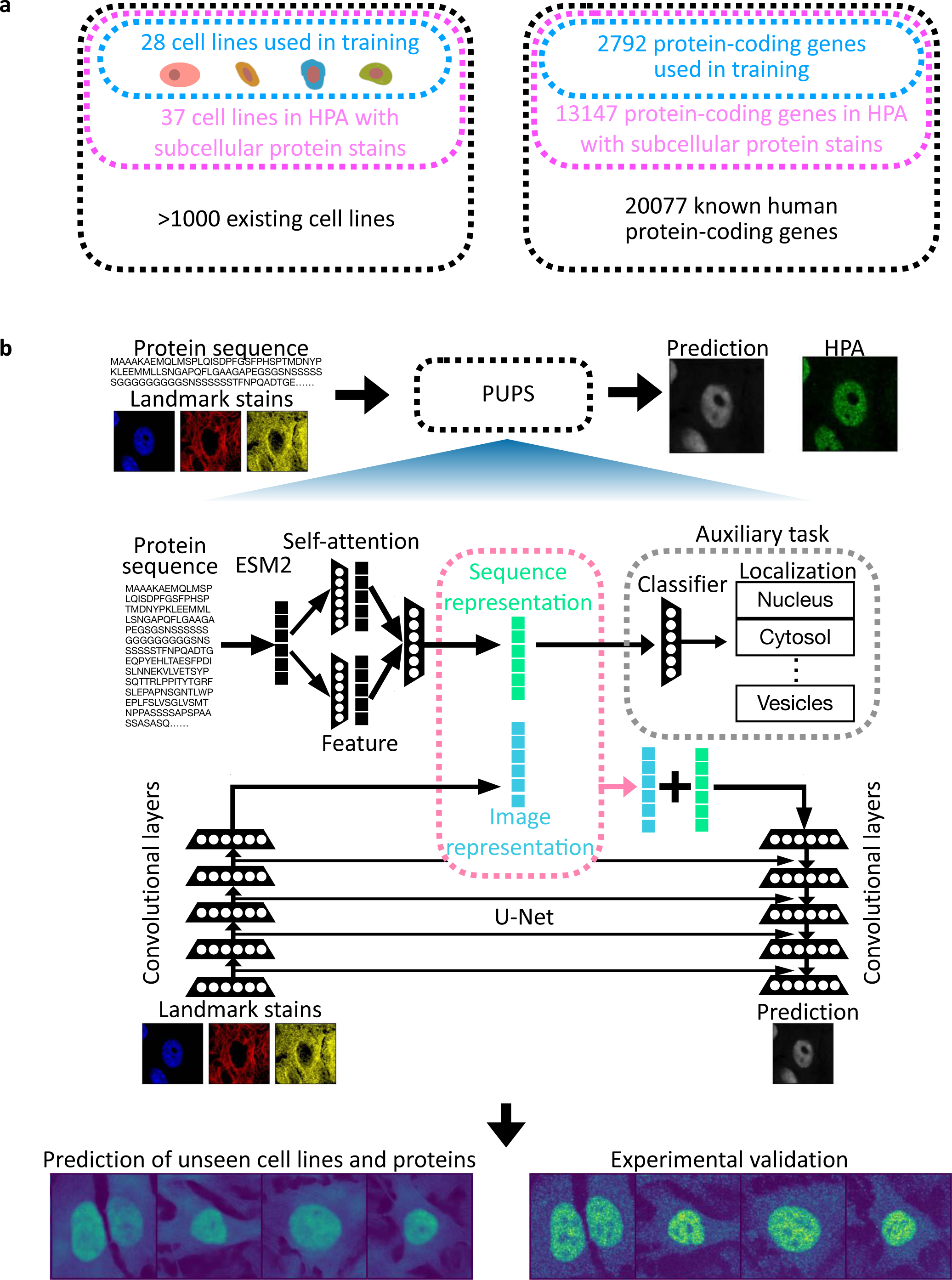
Our model enables the prediction of subcellular localization of unseen proteins in unseen cell lines. (a) There are 20,077 known human protein-coding genes (T2T-CHM13v2.0)^41^ and over 1,000 commercially available human cell lines. The Human Protein Atlas (HPA) provides the subcellular localization of proteins encoded by 13,147 genes, and each protein is only measured in up to three cell lines with a total of 37 cell lines in the entire dataset. We test the ability of our model to generalize to unseen proteins and cell lines by using 28 cell lines and 2,792 genes in training our model. (b) Our method predicts the subcellular localization of a protein in a given cell using both the protein sequence and the landmark stains of the cell. The landmark stains consist of the nuclear, microtubule, and endoplasmic reticulum (ER) stains. The protein sequence representation is learned using a pre-trained language model, ESM-2^7^, followed by a light attention layer (Methods). An auxiliary protein localization classification task using the sequence representation is trained jointly. The sequence representation is concatenated with the image representation of the landmark stains, learned using convolutional neural network layers, to output the protein image prediction (Methods). The use of both protein sequence and cellular landmark stains enables PUPS to predict the protein images of cells when both the protein and the cell line are not used in training the model. We performed validation experiments on cell lines and proteins not seen by our model, demonstrating that our model can be extended to experiments beyond HPA. Four examples of PSME3IP1 (Holdout 2) stained in HeLa cells (Holdout 1 and 2) are shown with both our model predictions and the images obtained from the validation experiments. All cell crops are 50μm x 50μm.

**Figure 2.**
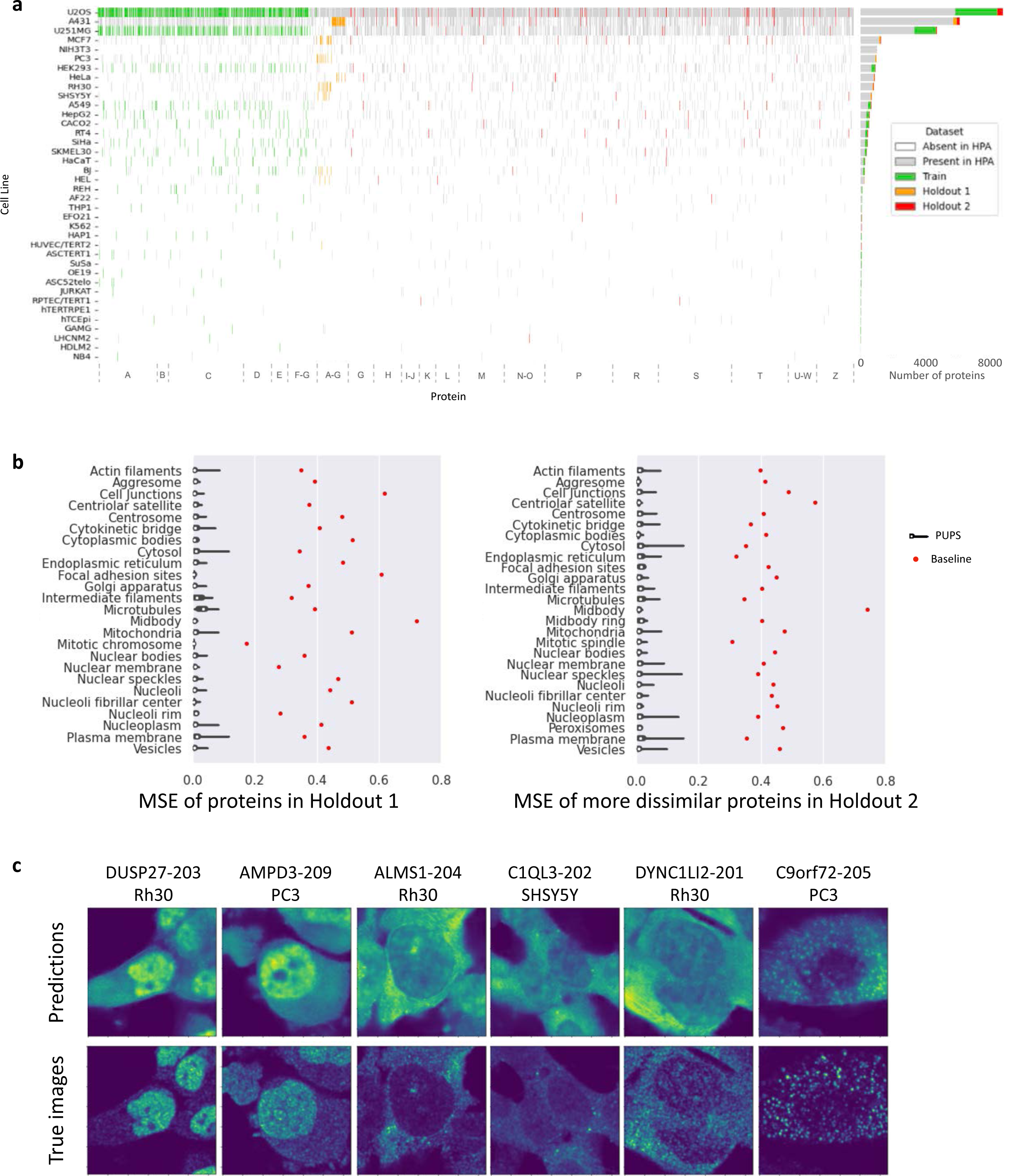
Our neural network based model learns a representation of both, protein sequence and cellular landmark stains, for accurate single-cell level prediction of unseen proteins in unseen cell lines. (a) Each protein in the HPA is measured in up to three cell lines in each version of HPA^3^. We aggregated multiple HPA versions to gather a more complete dataset (Methods). We alphabetically ordered all 13K proteins in the columns and indicated if a particular protein and cell line pair is in the training set (green), holdout set 1 (orange), holdout set 2 (red; consisting of more dissimilar proteins including unseen protein families), present in HPA but not used by our model (gray), or absent in HPA (white). (b) Mean-squared error (MSE) losses compared to the ground truth images in HPA are shown for held-out proteins and cell lines. The distribution of MSE losses are separately plotted for proteins annotated with different organelle localizations. We separate the held-out proteins into two groups with different degrees of similarity compared to the proteins used in training (Methods, Figure 2a). The left panel contains the test set of Holdout 1 with proteins that are more similar to the proteins used in training including proteins in the same protein family as the proteins used in training. The right panel contains Holdout 2 with proteins that are less similar to the proteins used in training including proteins from unseen protein families. The red dots indicate the average MSE loss of baseline predictions, obtained for each single cell with homogenous protein expression inside the cell. (c) Our model predictions and the real images from HPA are plotted for 6 randomly selected cells. All proteins and cell lines used in this plot are held-out from model training (Figure 2a, Holdout 1, Methods). All cell crops are 50μm x 50μm.

## Results

### 1. PUPS combines protein language and image inpainting models to predict protein subcellular localization

While large-scale atlases have been collected to annotate protein localization in various cell lines^3,4^, computational models are needed to predict protein localization for the thousands of proteins and cell lines not yet measured by the atlases. Our model, PUPS, consists of two major components, one for learning a sequence representation from the amino acid sequence of a protein and the other for learning an image representation from the landmark stains of a target cell (Figure 1b, Supplementary Figure 1-3). The combination of protein sequence representation and image representation is then used to predict the subcellular localization of the protein in the target cell. The use of both, protein sequence and cellular image, enables generalizing to unseen proteins and cell lines as well as capturing protein localization variability across cell lines and across single cells within a cell line.

The protein sequence representation from the amino acid sequence of a protein is learned by utilizing a pretrained ESM-2 language model, which has been shown to learn informative sequence features that lead to accurate protein structure prediction^7^ (Figure 1b, Supplementary Figure 2, Methods). An auxiliary pretext task is trained simultaneously to predict the subcellular compartment labels from the learned sequence representation. The image representation from the landmark stains of a cell is learned by a convolutional neural network^16^ (Methods, Figure 1b, Supplementary Figure 1). The cellular image and protein sequence representations from the two models are then combined to predict the protein image for a given cell through a convolutional network. All components of our model are trained simultaneously to minimize the classification loss of the pretext task and the difference between the predicted protein image and the experimentally measured protein image in the HPA.

We utilize the Cell Atlas in HPA^3^ to train our model and demonstrate that our model can accurately predict held-out protein images, even when the entire protein family is held-out from training. This shows that our model generalizes to proteins with less sequence homology or shared function. Towards this, we randomly held-out 9 cell lines and 10,355 proteins in the HPA for testing our model (Methods, Figure 2a). In particular, we computed the prediction loss of our model on two separate test sets, one with proteins that might be more similar to the proteins used in training (Holdout 1, Figure 2a) and the other with proteins that are from different protein families and more dissimilar in terms of sequence and function (Holdout 2, Figure 2a, Methods). A random baseline prediction is obtained for each single cell with the target protein homogeneously distributed inside the cell. For both test sets, our model accurately predicts the protein images for proteins in all cellular compartments (median mean-squared error loss of 0.00705 for Holdout 1 and 0.00960 for Holdout 2) and the prediction losses are much lower than the random baselines (median mean-squared error loss of 0.408 for Holdout 1 and 0.412 for Holdout 2, Figure 2b and 2c). Given the overlap in fluorescent spectra of the landmark stains and the target protein stain^15^ (Supplementary Figure 4a), evaluating the model predictions on unstained proteins is essential for ensuring that the model prediction does not rely on protein fluorescence bleed-through in the landmark images (Supplementary Figure 4b).

### 2. PUPS learns cellular context from the landmark stains to accurately predict differences in protein localization across cell lines

While protein localization is tightly coupled to its function, variability of protein localization across different cell types or cell lines has not been extensively studied. A previous study suggests that proteins with variable localization across cell lines might be more likely to be associated with carcinogenesis^17^. However, an extensive analysis of localization variability across cell lines is difficult given that HPA only measures up to three different cell lines for each protein, and the available cell lines are different between different proteins^3^. Thus, accurate prediction of protein localization in unseen cell lines could potentially lead to novel insights into the association between cell line variability in localization and different diseases.

To evaluate the performance of PUPS in quantifying the variability of protein localization across cell lines, we quantified the proportion of each protein localized in the nucleus, since nucleus and cytosol are the cellular compartments with the highest variation in protein localization^3^. We used the nuclear segmentation mask obtained from the nucleus image of each single cell to compute the proportion of each protein within the nuclear mask with respect to the total protein expression in the cell (Methods, Figure 3a). The intra-nuclear proportion of held-out proteins in predicted single-cell protein images is highly correlated with the real intra-nuclear proportion computed from the HPA images (Pearson correlation = 0.794 for Holdout 1 and 0.878 for Holdout 2, Figure 3a, Supplementary Figure 5a).

**Figure 3.**
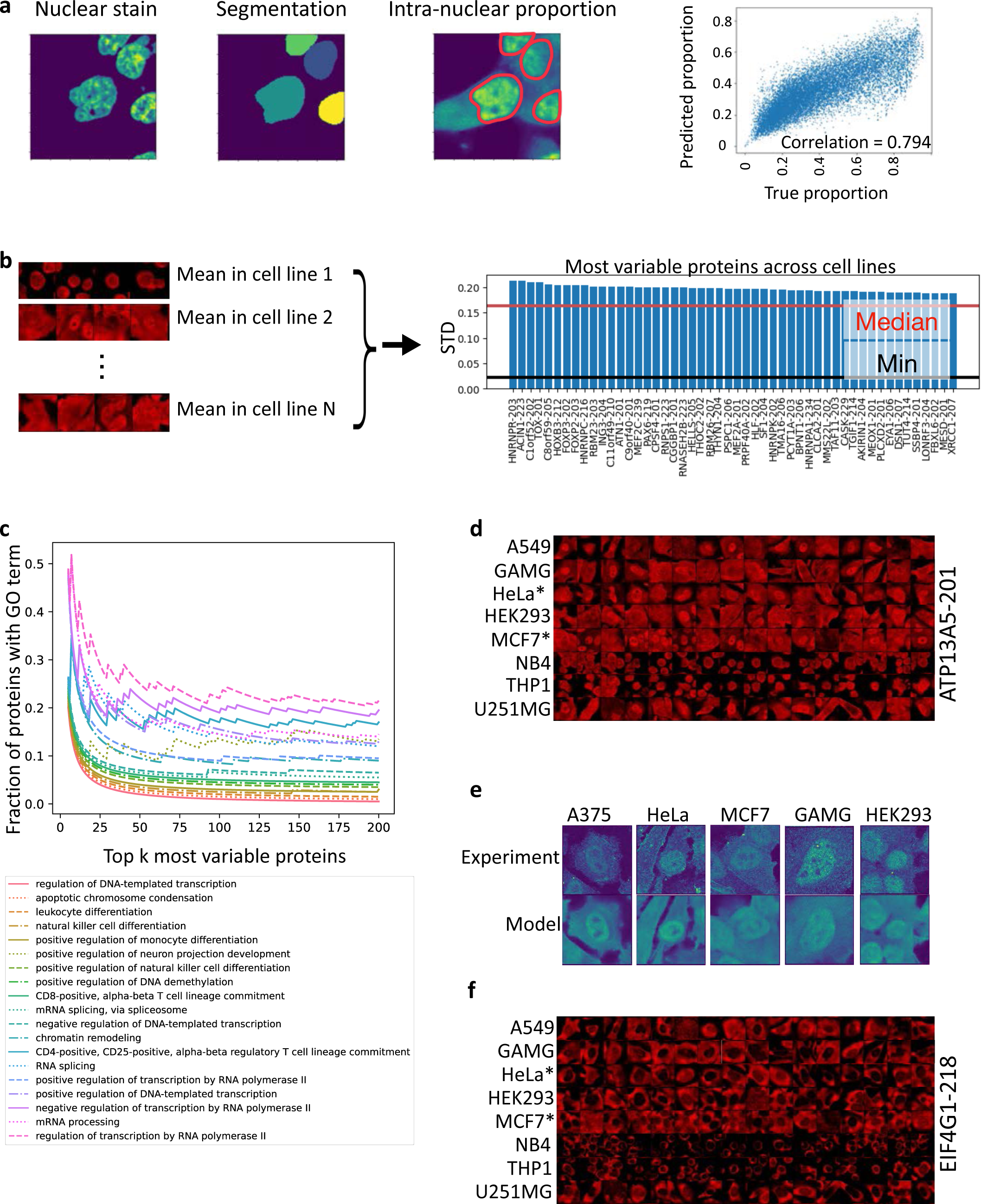
Our model learns cellular contexts from landmark stains to accurately predict differences in protein localization across cell lines. (a) Intra-nuclear proportion of a particular protein in a cell is quantified using the nuclear segmentation of the cell given its nuclear stain. The intra-nuclear proportion is computed as the sum of all protein pixels overlapping with the nuclear segmentation mask normalized by the sum of all protein pixels in the image. We plotted a scatter plot of all held-out cells (Holdout 2) comparing the intra-nuclear proportion computed using the predicted protein images to the proportion computed using real images from HPA. The Pearson correlation coefficient between the predicted intra-nuclear proportion and the experimentally measured proportion is 0.794. All cell lines and proteins used for plotting are not used in training the model, including proteins from unseen protein families that are not used in training. (b) We computed the variability of the intra-nuclear proportion of a particular protein across cell lines. Standard deviation (STD) of the mean intra-nuclear proportion across a total of 8 cell lines across both Holdout 1 (8 cell lines) and 2 (6 cell lines) are computed for all proteins (Methods, Supplementary Figure 5b). The proteins are ranked by their STD across the 6 cell lines in Holdout 2 and the most variable proteins are plotted (right figure). The red line indicates the median STD across all proteins. The black line indicates the minimum STD across all proteins. (c) We plotted the gene ontology (GO) terms of the top k most variable proteins in Holdout 2 for k ranging from 5 to 200. For each value of k, we computed the fraction of genes annotated with each biological process term in GO. The most frequently occurring terms are indicated in the legend, from the least frequent to the most frequent. In order to visually separate the GO terms, each term, from the least frequent (ranking = 0) to the most frequent (ranking = 18), is added with 0.005 * ranking of the term. Supplementary Figure 5d shows the plot without the additional values added. (d) For the 8 cell lines used in (b) and Supplementary Figure 5, we plotted the predicted images of one of the most variable proteins, ATP13A5 (in the training set), for 16 randomly selected cells. *: Cell lines are held-out from training the model. All cell crops are 50μm x 50μm. (e) Experimental validation of ATP13A5 (in the training set) in 5 cell lines. A375 is not in HPA and thus not plotted in d or f. The top row contains the real images of ATP13A5 from the validation experiment and the bottom row contains predicted images from our model. All cell crops are 50μm x 50μm. (f) For the 8 cell lines used in (b) and Supplementary Figure 5, we plotted the predicted images of the least variable proteins, EIF4G1 (in the training set), for 16 randomly selected cells. *: Cell lines are held-out from training the model. All cell crops are 50μm x 50μm.

Given the accurate prediction of intra-nuclear proportion on a single-cell level, we used the protein images predicted by our model to compute the standard deviation of each protein across cell lines and validate these experimentally (Methods, Figure 3b, Supplementary Figure 5b). While our predictions contain pairs of proteins and cell lines not measured in the HPA, experimental validation confirms that our prediction accurately captures the variability across cell lines for one of the most variable proteins, ATP13A5 (Figure 3d and 3e). Our analysis of the most variable proteins across all cell lines shows that the most variable proteins are associated with transcription, cell differentiation, and chromatin regulation, whereas the least variable proteins across cell lines tend to localize in the cytosol but not in the nucleus (Figure 3c, 3d, 3f, and Supplementary figure 5c-i). This is in contrast to a previous study using mass spectrometry that shows nuclear proteins have more stable overall expression level between cell lines than cytoplasmic proteins^5^, possibly indicating that the subcellular distribution of proteins can vary significantly while having stable overall expression level.

### 3. PUPS learns cellular context from the landmark stains to accurately predict variability in protein localization across single cells within a cell type

While localization of a particular protein in multiple subcellular compartments has been observed^3,5^, such variability is not well quantified and studied across proteins in different cell lines. Besides the potential association of single-cell variability with cell cycle^3^, it is also not well understood to what extent the single-cell variability is stochastic. A model that can predict protein localization in single cells within a cell line is needed to facilitate the understanding of single-cell variability in protein localization. While there have been limited studies in single-cell protein localization prediction, previous works have shown that cell/nucleus morphology is tightly coupled to and predictive of gene expression.^18–20^ Thus, we hypothesized that the image representation of PUPS derived from cellular landmark stains could also enable the prediction of protein localization at a single-cell level and provide an opportunity to quantify single-cell variability for any pair of protein and cell line. Additionally, whether the single-cell variability of protein localization is predictable from the landmark stains could also suggest the degree of stochasticity in protein localization.

For each pair of held-out cell lines and proteins, we computed the variance of the intra-nuclear proportion of the protein across all single cells within each cell line (Figure 4a). The predicted ranking of single-cell variability for each protein and cell line pair shows high consensus with the real ranking (Figure 4b and Supplementary Figure 6a). The predicted distribution of intra-nuclear protein proportion across single cells for the most variable and the least variable proteins highly correlates with the real distribution; and the variability in intra-nuclear proportion is not a result of prediction errors (Figure 4c, 4e, and Supplementary Figure 6b and 6c). This indicates that high variability in protein localization across single cells is not only due to stochasticity but is predictable from cell morphology. Gene ontology (GO) analysis shows that the highly variable proteins are associated with cell division, transcription, double-strand break repair, and apoptosis, which confirms that cell cycle related processes could explain some of the single-cell variability (Figure 4d and and Supplementary Figure 6e).

**Figure 4.**
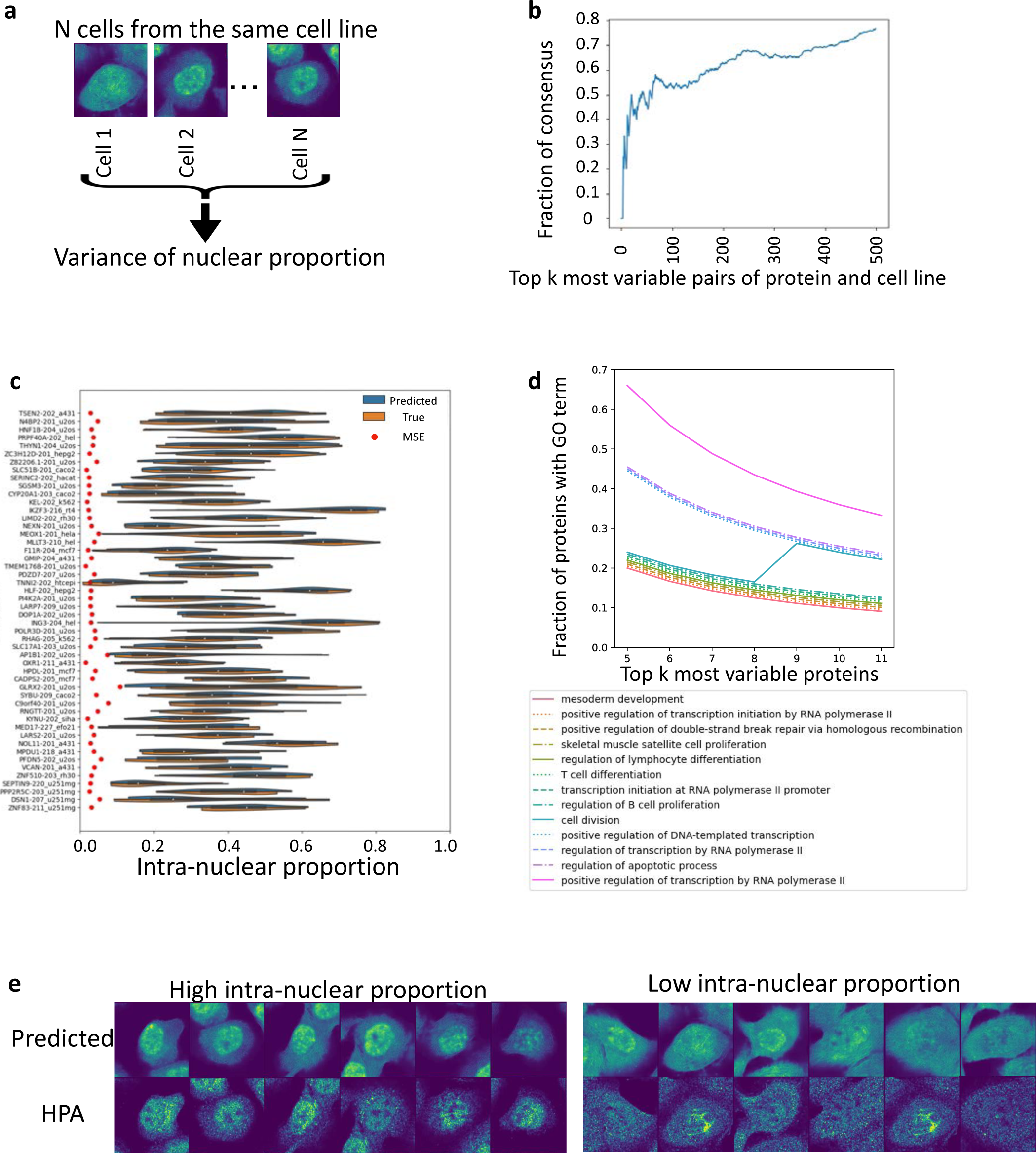
Our model can predict variability in protein localization across single cells within a cell line. (a) For each protein-cell line pair, we computed the variance of intra-nuclear proportion of the protein across single cells in the cell line. (b) Protein-cell line pairs in Holdout 2 are ranked by the variance of intra-nuclear proportion. We compared this ranking computed with predicted protein images using held-out protein-cell line pairs to the ranking computed with real protein-cell line pairs available in the HPA. The fraction of protein-cell line pairs that overlap between the two rankings is plotted for the top k most variable pairs for k ranging from 1 to 500. All cell lines and proteins used for plotting are not used in training the model and include proteins from unseen protein families that are not used in training. (c) For the top 50 most variable protein-cell line pairs in Holdout 2 that overlap between the predicted and real HPA images, we plotted the distribution of intra-nuclear proportion for both the predicted and the real HPA images. Red dots indicate the average MSE loss of the predicted images compared to the real HPA images, demonstrating that the variability in intra-nuclear proportion is not a result of prediction error. (d) We plotted the gene ontology (GO) terms of the top k most variable proteins in Holdout 2 for k ranging from 5 to 11. For each value of k, we computed the fraction of genes annotated with each biological process term in GO. The most frequently occurring terms are indicated in the legend, from the least frequent to the most frequent. In order to visually separate the GO terms, each term, from the least frequent (ranking = 0) to the most frequent (ranking = 12), is added with 0.005 * ranking of the term. Supplementary Figure 6d shows the plot without the additional values added. (e) We plotted examples of predicted and real HPA protein images for one of the most variable protein-cell line pairs, protein TSEN2 in cell line A431. TSEN2 is in Holdout 2 and A431 is in both Holdout 1 and 2 (Figure 2a, Methods). All single cells are divided by their intra-nuclear protein proportions into three groups of equal percentile intervals; 6 randomly selected cells in the groups with the highest and lowest intra-nuclear proportions are plotted for both the predicted and real HPA protein images. All cell crops are 50μm x 50μm.

### 4. PUPS enables the prediction of protein subcellular localization in experiments beyond the HPA

Given the high variability of protein localization, it is important for any prediction model to be able to generalize to new experiments to predict protein localization in specific cellular contexts. We selected 9 proteins for experimental validation in 5 cell lines based on our analysis of variability in intra-nuclear proportions (Figure 5a and Supplementary Figure 7-11). ATP13A5, CHID1, COPA, MESD, and RBM23 were selected as the most variable proteins across cell lines that each have a distinct GO term (Methods, Figure 3c, Supplementary Figure 5e-h). DDIT3 and N4BP2 are the most variable proteins across single cells within a cell line (Methods, Figure 4c, Supplementary Figure 6b). EIF4G1 and PSME3IP1 are the least variable proteins across cell lines with EIF4G1 expected to be mainly outside the nucleus and PSME3IP1 expected to be mainly inside the nucleus (Figure 3f, Supplementary Figure 5i and 5j). We validated the localization of these proteins in 5 cell lines (A375, HeLa, MCF7, GAMG, HEK293), all of which except A375 are also contained in HPA. The protein images predicted by our model are visually similar to the experimentally measured images, including for A375, which is not present in the HPA (Figure 5a, Supplementary Figure 7-11) and can also be used for quantitative assessment of localization. The intra-nuclear protein proportion of each single cell computed using the predicted protein images is strongly correlated with the proportion computed from the experimentally measured images (Pearson correlation = 0.767, Figure 5b). Using the predicted intra-nuclear proportions to quantify variability in protein localization across cell lines (Figure 5d) and across single cells within a cell line (Figure 5c) both show high consensus with the real variability. Our results demonstrate that PUPS can be used to quantitatively predict the localization of proteins that were not previously experimentally measured or used in the training atlas.

**Figure 5.**
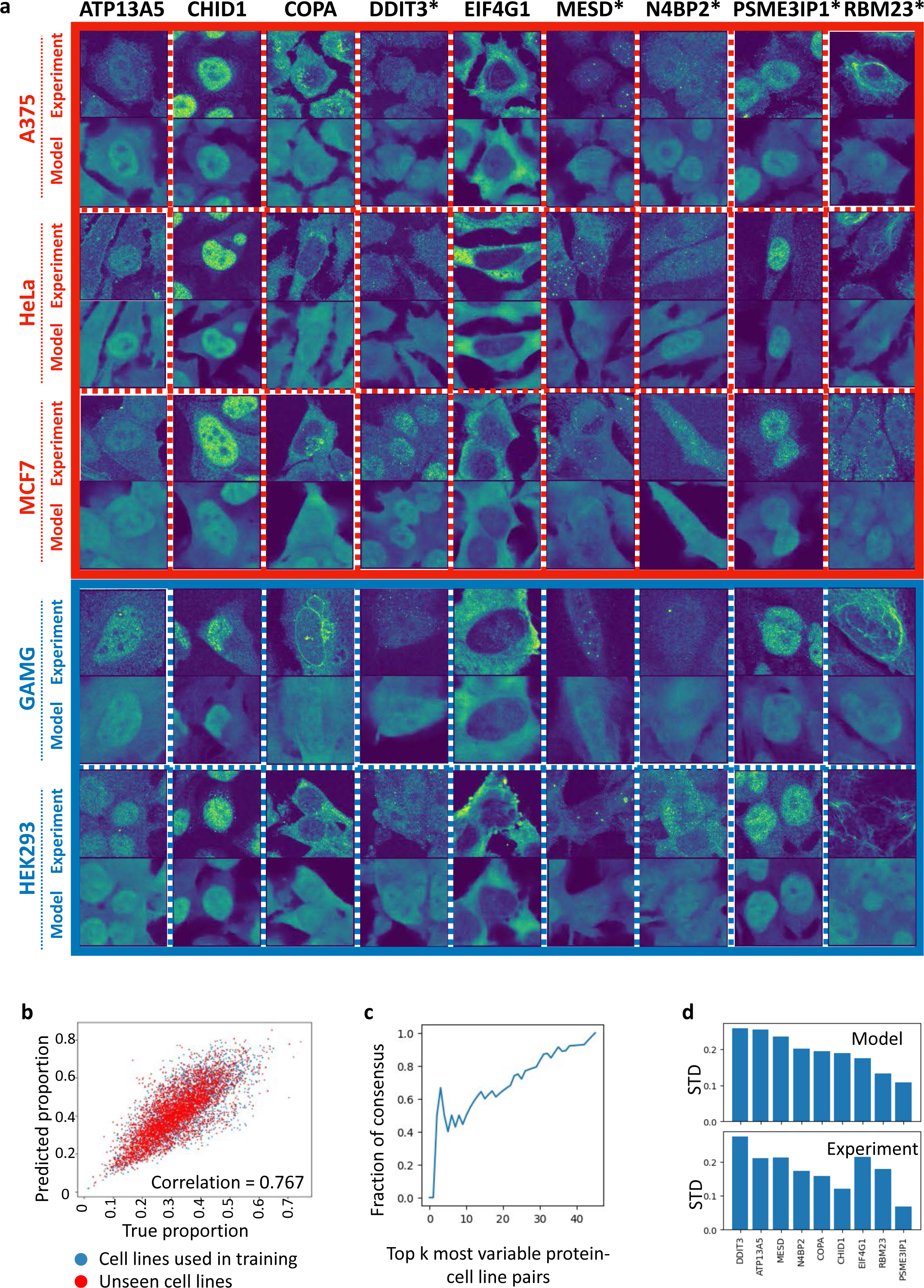
Our model enables the prediction of protein subcellular localization in experiments beyond the HPA. (a) We performed imaging experiments on 5 cell lines (rows) and 9 proteins (columns) (Methods); 3 out of the 5 cell lines are not used in training the model (red box); proteins not used in training the model are marked by asterisks. The proteins were selected for experimental validation based on their variability in intra-nuclear proportions. One example of a predicted protein image and the corresponding real image from our experiment is plotted for each cell line and protein pair. All cell crops are 50μm x 50μm. (b) We plotted a scatter plot of all cells comparing the intra-nuclear proportion computed using the predicted protein images to the proportion computed using real images from the experiments. Cells from the two cell lines used in training the model are colored in blue. Cells from the held-out cell lines not used in training the model are colored in red. The Pearson correlation coefficient between the predicted intra-nuclear proportion and the experimentally measured proportion is 0.767. (c) All protein-cell line pairs are ranked by the variance of intra-nuclear proportion. We compared this ranking computed with predicted protein images to the ranking computed with real protein images measured in the experiments. The fraction of protein-cell line pairs that overlap between the two rankings is plotted for the top k most variable pairs of proteins and cell lines for k ranging from 1 to 45. (d) Top panel: Standard deviation (STD) of the mean intra-nuclear proportion across the 5 measured cell lines is computed for all 9 proteins. The proteins are ranked by their STD across the cell lines. Bottom panel: Standard deviations computed from real images obtained from the experiments are plotted.

### 5. PUPS learns meaningful protein and cell representations

We demonstrate that the ability of our model to predict protein localization in unseen proteins and cell lines results from learning meaningful representations of protein sequences and cellular landmark images. We plotted the protein sequence representation of 40,622 proteoforms corresponding to 12,614 genes. Proteins with similar localization annotations tend to have similar sequence representations (Figure 6a and Supplementary Figure 12). To further demonstrate that our model recognizes meaningful protein sequence patterns to predict localization, we computed the importance of each amino acid residue in a particular protein to the prediction of each compartment label using the Positional Shapley method^21–23^ (Figure 6b and Methods). We visualized the amino acid residue importance of a held-out mitochondrial protein, AARS2, with an N-terminal mitochondrial transit peptide annotated by UniProt^24^ (Figure 6b). Our model indeed identified the N-terminal residues of AARS2 as predictive of mitochondrial and extra-nuclear localization (Figure 6b). Furthermore, we visualized the amino acid residue importance of a held-out nucleoplasmic protein, DDIT3. Our model correctly predicted its compartment label and identified the basic leucine zipper domain, which is known to contain a nuclear localization signal (NLS)^25,26^, as important for its prediction (Figure 6b).

**Figure 6.**
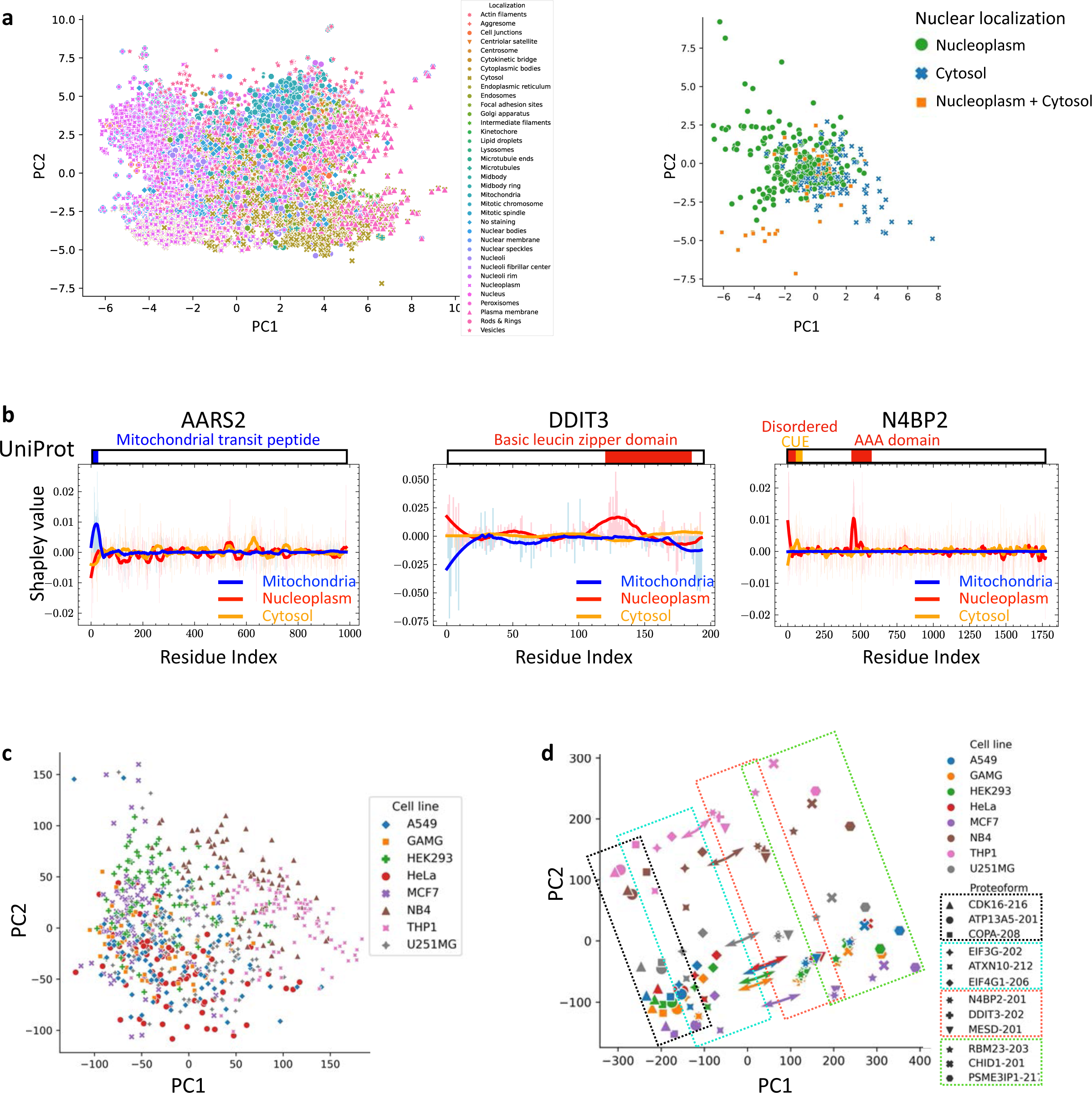
PUPS learns meaningful protein and cell representations. (a) The sequence representations (Figure 1b) of 40,622 proteoforms corresponding to 12,614 genes are computed. Left panel is a visualization of all 40,622 proteoforms with each protein colored by its localization annotation in the HPA. Right panel is a visualization of a subset of proteins in the test set of Holdout 1 which localize in nucleoplasm, cytosol, or both. (b) We computed the importance of each amino acid residue in a particular protein to the prediction of each compartment label using Positional Shapley values^21–23^ (Figure 1b auxiliary task; Methods). An example of residue importance in three proteins to the prediction of mitochondria, nucleoplasm, and cytosol labels are shown (left and middle panels). We compare the residue importance derived from our model to the UniProt features^24^. (c) The image representations of the landmark stains in the test set of Holdout 1 (Figure 1b) are plotted for the 8 cell lines used for computing protein localization variability in Figure 3. (d) Joint representations of proteins and landmark images are the output of the first convolutional layer after the concatenation of sequence and image representations (Figure 1b). The mean representation of each protein-cell line pair is plotted. Arrows of each cell line start from the centroids of the joint representation of the particular cell line and point towards the direction of ATP13A5 and CHID1 of the same cell line. Proteoforms are grouped based on their relative locations within each cell line, e.g. CDK16, ATP13A5, and COPA have the smallest PC1 and PC2 values within each cell line.

Interestingly, phosphorylation within the NLS region of DDIT3 is known to alter its intra-nuclear proportion^26^ and our model is able to predict the intra-nuclear proportion of DDIT3 in different cells using only the cellular landmark stains (Figure 5a, Supplementary Figure 8). We further visualized the predictive residues of N4BP2, a held-out protein for which little was known regarding its subcellular localization and which was predicted by PUPS to have a highly variable intra-nuclear proportion across single cells (Figure 5a and 6b). Our analysis shows that the N-terminal disordered domain and the AAA domain are predictive of N4BP2’s localization to nucleoplasm, while its ubiquitin-binding CUE domain is predictive of localization to the cytosol and is consistent with the observation that CUE domain promotes endosomal (i.e. cytosolic) localization in other proteins^27^. This could explain N4BP2’s predicted variability in nuclear localization, which is also consistent with the reported role of CUE domain in ubiquitin binding that might alter subcellular localization^28^.

In addition to recognizing meaningful protein sequence motifs, we further show that PUPS learns meaningful representations of single cells from cellular landmark stains. Towards this, we visualized the image representations of single cells learned from landmark stains and found that single cells of the same cell line have similar image representations, even though the cell line labels are not input to our model (Figure 6c and Supplementary Figure 13). The joint representation of proteins and landmark images of cells retain the separation between both cell lines and proteins, while the different proteins within each cell line are ordered similarly across different cell lines (Figure 6d). Given the centroid of each cell line in the joint representation space, the vector from the centroid to a particular protein is mostly parallel across all cell lines (Figure 6d), i.e., predicting the image of a particular protein requires moving in the representation space in the same direction given the sequence representation regardless of the cell line. This explains the capabilities of PUPS to generalize to unseen proteins and cell lines by learning meaningful protein and cell image representations.

## Discussion

The availability of protein atlases provides important insights into the subcellular localization of proteins and the variability of protein localization across cell lines and single cells^3–5^. However, a significant number of proteins and cell lines are not measured in current protein atlases (Figure 1a and 2a). Additionally, the variability of protein localization in single cells suggests that the localization of proteins in experiments outside of the existing atlases could be different from the localization observed in the existing atlases. New computational methods are needed to predict protein localization for the thousands of proteins and cell lines not yet measured in atlases.

Recent advances in machine learning have aided the use of protein atlases by predicting missing data^7,8,15^ as well as providing automatic annotation of protein properties^13,14^. Existing computational methods cannot make predictions of protein localization in single cells for protein-cell line pairs that are not measured in the training data^13–15^. We presented a novel machine learning model, PUPS, that overcomes this limitation by learning representations for both protein sequences and cellular landmark stains of single cells to predict the localization of unseen proteins in single cells from unseen cell lines. We demonstrated that PUPS generates accurate predictions for held-out proteins and cell lines, including when the entire protein family is held-out from training. Through experimental validation we showed that the protein localization predictions of PUPS generalize to new experiments outside of HPA.

While variability in protein localization has been observed across cell lines and single cells within a cell line^3,5^, a comprehensive comparison across proteins and cell lines has been difficult given that HPA only measures up to three different cell lines for each protein and the available cell lines are different between different proteins^3^. Our model enables an accurate quantification of variability in protein localization across any protein-cell line pair, including pairs not measured in the HPA. Our analysis of the proteins predicted to be most variable in their localization across cell lines indicates that these proteins are associated with transcription, cell differentiation, and chromatin regulation, while the proteins predicted to be most variable across single cells within a cell line are associated with cell division, transcription, double-strand break repair, and apoptosis. We also validated experimentally the predicted variability of protein localization across cell lines and across single cells of the same cell line. We showed that PUPS extracts protein sequence features that are known to be relevant for localization (such as N-terminal residues, nuclear localization signals, AAA domains, and CUE domains) and learns meaningful representations of proteins and cell lines. This may explain PUPS’ ability to generalize to unseen proteins and cell lines.

The ability of PUPS to predict images of unmeasured proteins for experiments outside of the HPA used for training opens avenues to virtually stain cells for any protein using only the cellular landmark stains. This could also overcome experimental limitations in the number of proteins that can be measured simultaneously in multiplexing experiments^6^. Virtual staining of cells by predicting images of any protein could enable large-scale screening of proteins for the identification of disease biomarkers as well as the prediction of genetic or drug perturbation effects. The current model could be further extended to incorporate available protein images together with the cellular landmark stains to potentially improve the prediction of unmeasured proteins. Similar to the L1000 platform for gene expression measurement that only measures a small subset of the full transcriptome^29^, a careful selection of which proteins to measure combined with a prediction model for unmeasured proteins could allow us to gain more insights into protein subcellular localization and protein-protein interaction beyond the number of proteins that can be simultaneously measured in the same samples. While we showed that PUPS provides accurate predictions in cell lines, transcriptional differences have been reported between cell lines and the corresponding tissues of origin^30,31^. Analyzing and extending our model to predict protein subcellular localization in the tissue context is an interesting avenue for future work.

## Methods

### 1. Model architecture and training

PUPS consists of two major components, one for learning a sequence representation from the amino acid sequence of a protein and the other for learning an image representation from the landmark stains of a target cell, namely the nucleus, microtubule, and endoplasmic reticulum (ER) stains (Figure 1b). Two tasks are trained jointly using the sequence and the image representations - the main protein image prediction task that uses both representations and an auxiliary pretext task that classifies the protein localization using the protein sequence representation. All parameters are optimized using the Adam optimizer^32^ with a learning rate of 1e-4. The two model components are further described below. We plot the training and evaluation losses for 22 epochs with 64K cells used for training and 6.4K cells used for evaluation per epoch (Supplementary Figure 3). All figures and supplementary figures are generated with the model trained for two epochs.

#### 1.1. Protein language model

Our model learns sequence representations by using a language model, self-attention layer, and an auxiliary pretext task that classifies the protein localization given the learned sequence representation. An initial representation of a particular proteoform is obtained by inputting the sequence of the first 2000 amino acids from the N-terminus to the pretrained ESM-2 model to obtain a 1280-dimensional representation for each amino acid residue^7^.

Proteoforms with less than 2000 residues are padded with zeros. This sequence length cutoff was implemented to avoid biasing prediction towards the few proteins with sequence lengths up to tens of thousands of residues. To adapt the ESM-2 representation to predict protein localization, a light attention layer^33^ using separable convolutions^16^, which was found to perform well in protein sequence to localization predictions^13^, is applied to the ESM-2 representation to obtain a 300-dimensional sequence representation. This protein sequence representation is used for both the auxiliary pretext task of predicting localization labels and for protein image predictions when combined with the image representation. The pretext task inputs the protein sequence representation to a single fully connected neural network layer to output a 29-dimensional vector representing the probability of having each of the 29 subcellular compartment localization labels. The pretext task output is compared with the HPA annotated protein compartments using a binary cross entropy loss with sigmoid activation.

#### 1.2. Image inpainting model

The image input of each cell consists of three image channels, for nucleus, microtubule, and ER stains, and has a dimension of 3x128x128 centered around the centroid of the cell nucleus. Our model encodes the image input by 5 separable convolutional layers^16^ to a final dimension of 16x16x512 (Figure 1b). Each convolutional layer is followed by leaky ReLu activation^34^, batch normalization, and a 2D max-pooling layer with a kernel size of 2. The protein sequence representation is concatenated to all spatial dimensions of the cellular image representation, which is then input to the image decoder. The decoder consists of 5 separable convolutional layers and generates a 1x128x128 image output, which is the prediction of the protein image for the given cell. Skip connections, similar to U-Net for image segmentation^35^, are added between the encoding layers for generating image representations from landmark stains and the decoding layers at the same depth for generating protein image predictions (Figure 1b). Our model is trained to minimize the difference between the predicted protein images and the experimentally measured protein images using mean-squared error loss.

### 2. Data preprocessing

#### 2.1 Subcellular localization data from the HPA

We aggregated the HPA data from versions 16 (the first release of the Cell Atlas that measures the subcellular localization of proteins^3^) to 22 to gather as much data as possible for each protein, in terms of the number of cells measured, the number of cell lines stained for that protein, and the different antibodies potentially targeting different proteoforms of the protein.

This could provide a more comprehensive representation of the dependence of protein localization on cell lines and protein sequences. For example, one of the most highly variable proteins, TSPAN6 (Figure 4c), is stained in different cell lines in HPA version 16 and 22 and has different localization in the different cell lines. Since HPA uses polyclonal antibodies targeting multiple protein isoforms^3^, we also incorporated all the targeting proteoform sequences for each protein and the corresponding cellular compartment localization labels, when training or evaluating our model. The sequence of each proteoform is obtained from Ensembl API v103^36^.

#### 2.2 Preprocessing of cellular images

We followed the image preprocessing procedures in a previous work using HPA images^15^. Each image is downsampled 4 times to 0.32 μm/pixel. We segmented the nucleus channel by first obtaining a per-pixel intensity threshold using the Otsu method^37^ implemented in the scikit-image package^38^. Gaussian blurring (sigma = 5) is applied to the nucleus channel and the resulting image is filtered using the Otsu threshold. Small holes in the image are removed using the remove_small_holes function with the area threshold set to 300. The images are then binarized as nucleus masks and nuclei smaller than 100 pixels or at the image borders are removed. The centroid of each nucleus is obtained from the binary mask and used to define a 128 pixel x 128 pixel crop used for each cell. The intensity in each channel of a particular crop is separately rescaled to between 0 and 1. All landmark stain channels are additionally filtered for low intensity pixels, with a threshold of 0.19 to remove bleed-through from the targeting protein channel due to the overlapping spectra (Supplementary Figure 4).

Images from the validation experiments were processed similarly to the HPA images with additional steps to adjust for the fluorescent background. All images are rescaled to 0.32 μm/pixel to match the resolution used for the HPA images. The 3D stack of each image is first projected to 2D using the maximum intensity and the pixel intensity is capped at 99.92 percentile to reduce the impacts from outlier pixels with very high intensities. The images are then rescaled to between 0 and 1. Centroids of nuclei are identified using the same procedure described for the HPA images and cell crops are obtained using the centroids.

#### 2.3 Training and held-out datasets

The training dataset (340,553 cells) and the holdout dataset 1 (36,552 data points) were constructed from cell crops of 9,472 proteoforms (8,086 in the training set) corresponding to 3,312 genes (2,801 genes in the training set) with names beginning with letters between A and G across all 37 cell lines present in the HPA. The holdout dataset 1 was further randomly divided into an evaluation set and a test set, with 11,050 and 25,502 cells respectively. The holdout dataset 2 (24,007 data points) was constructed from cell crops of 556 proteoforms corresponding to 515 genes with names beginning with all letters of the alphabet. The alphabetical split of the training dataset and the holdout dataset 1 was taken so that the model’s ability to generalize to proteoforms from completely unseen gene families could be evaluated with the holdout dataset 2. The protein encoded by each gene is present in at most one dataset. The training dataset and the holdout dataset 1 have no overlapping cell lines. See Figure 2a for a visualization of the dataset split. 10 additional genes were also included in the training dataset outside of the alphabetical ordering, namely: IHO1, IMPAD1, INKA1, ISPD, ITPRID1, KIAA1211L, KIAA1324, LRATD1, SCYL3, TSPAN6. BJ cell line images were included in both the training and holdout 1 datasets. During the data pipeline development these were used as the first prototype targets and were thus included in the final dataset.

### 3. Protein localization variability

We used the pretrained StarDist model^39^ to obtain the nuclear segmentation mask of each cell using the nucleus image. The intra-nuclear proportion of a cell is computed as the proportion of each protein within the nuclear mask with respect to the total protein expression in the cell (Figure 3a). For calculating the variability of nuclear localization across cell lines (Figure 3b), the mean intra-nuclear proportion in each cell line is computed based on thresholded intra-nuclear proportion: 1) Cells with intra-nuclear proportion greater than 2/3 are labeled with 1; 2) Cells with intra-nuclear proportion less than 1/3 are labeled with -1; 3) all other cells are labeled with 0. The 8 cell lines in Figure 3 and Supplementary Figure 5 and 6 are selected based on both experimental availability and availability in the HPA. 4 of the 8 cell lines were selected for their availability for experimental validation (HeLa, MCF7, GAMG, HEK293). The remaining 4 cell lines were selected to cover a diverse range of tissue types as well as availability in HPA, e.g. U251MG is one of the cell lines with the most amount of data in HPA and NB4 is the cell line with the least amount of data (Figure 2a). NB4 and THP1 are not in Holdout 2 and are thus omitted from plotting. For calculating the variability of single cells in the same cell line, the intra-nuclear proportion of each cell is computed using the nuclear mask as described above.

#### 3.1 Gene ontology of most variable proteins

For both the most variable proteins across cell lines (Figure 3c) and the most variable proteins across single cells within a cell line (Figure 4d), the same procedure is used for computing the gene ontology (GO) terms that are most represented by the genes encoding the top k most variable proteins. For each value of k, the fraction of top k genes annotated with each GO term is calculated. The ranking of the GO terms in Figure 3c and 4d and Supplementary Figure 5c, 5d, 6d, and 6e are in terms of the maximum values along the curves.

### 4. Positional Shapley analysis of amino acid residue importance for localization prediction

We adapted the SHapley Additive exPlanations (SHAP) library^21^ to compute the importance of each amino acid residue, given its 1280 dimensional ESM-2 representation, for the prediction of the localization label (Figure 1b, auxiliary task). We used the Positional Shapley method from previous works^22,23^, based on the additive nature of the Shapley values, to swap the 1280-dimensional amino acid representation of a particular protein with the

1280-dimensional baseline feature vector. The feature baseline was constructed from the average of 300 proteoforms with sequence length equal to or greater than the target proteoform. A subset of amino acids was selected by the SHAP kernel explainer at each time to be swapped with the baseline features at the same locations along the sequence and the effect on the localization label prediction was recorded. This swapping was repeated for 10,000 times using the SHAP kernel explainer to generate positional Shapely scores which were then smoothed with the Savitzky-Golay filter of window length 51 and polynomial of order 2 (Figure 6b)^40^.

### 5. Experimental validation

#### 5.1 Chemicals

Chemicals were purchased from Sigma-Aldrich if not specified otherwise; no further purification was performed before usage.

#### 5.2 Tissue culture cell line cultivation

All cell lines were cultivated at 37°C in a 5% CO_2_ humidified incubator with medium supplied with 10% fetal bovine serum (FBS; Sigma-Aldrich, Cat# F4135), and 1% Pen Strep. Specifically, HeLa and HEK293 cells were cultivated in 1× DMEM with GlutaMax-1 (Gibco, Cat# 10569010), while MCF7, GAMG, and A375 cells were cultivated in RPMI 1640 (Gibco, Cat# 11835055).

#### 5.3 Immunofluorescence staining

For the immunofluorescence experiments, 96-well glass bottom plates (Greiner Bio-One, Cat# 655892) were coated with 12.5 µg/ml fibronectin (Sigma-Aldrich, Cat# F0895) diluted in 1× PBS for 1 h at room temperature (RT). Then, wells were washed with 1× PBS, and 14,000 cells were seeded in each well to grow overnight. On the second day, cells were washed with 1× PBS, then fixed in 4% paraformaldehyde (PFA) in 1× PBS for 15 min (200 μL/well), followed by permeabilization with 0.1% Triton X-100 in 1× PBS for 3×5 min. The cells were quickly rinsed with 1x PBS. Before immunostaining, the cells were incubated with a blocking buffer (4% FBS in 1× PBS) for 5 min. After another quick washing step with 1× PBS, the cells were incubated with the 1^st^ antibodies overnight at 4 °C.

The clone, host, and final concentration of each antibody used in the staining can be found in Supplementary Table 1, all wells were stained with the 1^st^ antibodies against a-Tubulin and calreticulin, but with different rabbit polyclonal antibodies from the Human Protein Atlas (HPA) purchased from Atlas Antibodies. On the next day, the cells were first washed with 1× PBS for 4×10 min, then incubated with the secondary fluorescent antibodies diluted with blocking buffer for 90 min at RT. Cells were subsequently counterstained with 4′,6-diamidino-2-phenylindole (DAPI) diluted in 1× PBS for 10 min, then washed with 4×10 min washes with 1× PBS, then kept in 1× PBS under 4 °C from light until imaging.

#### 5.4 Fluorescence image acquisition

Fluorescent images were acquired with an Andor Dragonfly 200 spinning disk confocal microscope equipped with a Nikon Plan Apo 60×/1.4 oil objective. Here, z-stacks of at least six field-of-views (FOVs) of 205.8×205.8 μm were acquired for each cell line and HPA antibody combination, with pixel size of 0.1005 μm/pixel. All cells from all FOVs obtained in the experiments were used for the evaluation of our model performance (Figure 5b-d).

## Data availability

All images from the Human Protein Atlas (HPA) can be downloaded from the HPA website: https://www.proteinatlas.org/humanproteome/subcellular. Images from the experimental validation experiments have been uploaded to the Google Drive folder for the review process and will be deposited to a public repository upon publication: https://drive.google.com/drive/folders/1olCw88ojd0yZTl7ivx2lSkkeQlvmI53t?usp=share_link.

## Code availability

The code is available at https://github.com/uhlerlab/PUPS

## Acknowledgement

X.Z. was supported by a fellowship from the Eric and Wendy Schmidt Center at the Broad Institute. Y.T. was supported by the NIH Graduate Research Fellowship Grant (2141064, 1745302). F.C. acknowledges support from the NHGRI (R01, R01HG010647), the Burroughs Wellcome Fund CASI award, the Searle Scholars Foundation, the Harvard Stem Cell Institute, and the Merkin Institute. C.U. was partially supported by NCCIH/NIH (1DP2AT012345), ONR (N00014-22-1-2116), and the United States Department of Energy (DE-SC0023187).

