## Supplementary files for "Prediction of protein subcellular localization in single cells"

**Supplementary Information:**
**Prediction of protein subcellular localization in single cells**

Xinyi Zhang[1,2,*], Yitong Tseo[3,*], Yunhao Bai[4], Fei Chen[4,5,#], Caroline Uhler[1,2,#]
1 Department of Electrical Engineering and Computer Science, Massachusetts Institute of Technology, U.S.A.
2 Eric and Wendy Schmidt Center, Broad Institute of MIT and Harvard, U.S.A.
3 Computational and Systems Biology Program, Massachusetts Institute of Technology, U.S.A.
4 Broad Institute of MIT and Harvard, U.S.A.
5 Department of Stem Cell and Regenerative Biology, Harvard University, U.S.A.

*: Equal contribution

**This PDF file includes:**

Supplementary Figures 1-13

Supplementary Table 1

Supplementary References

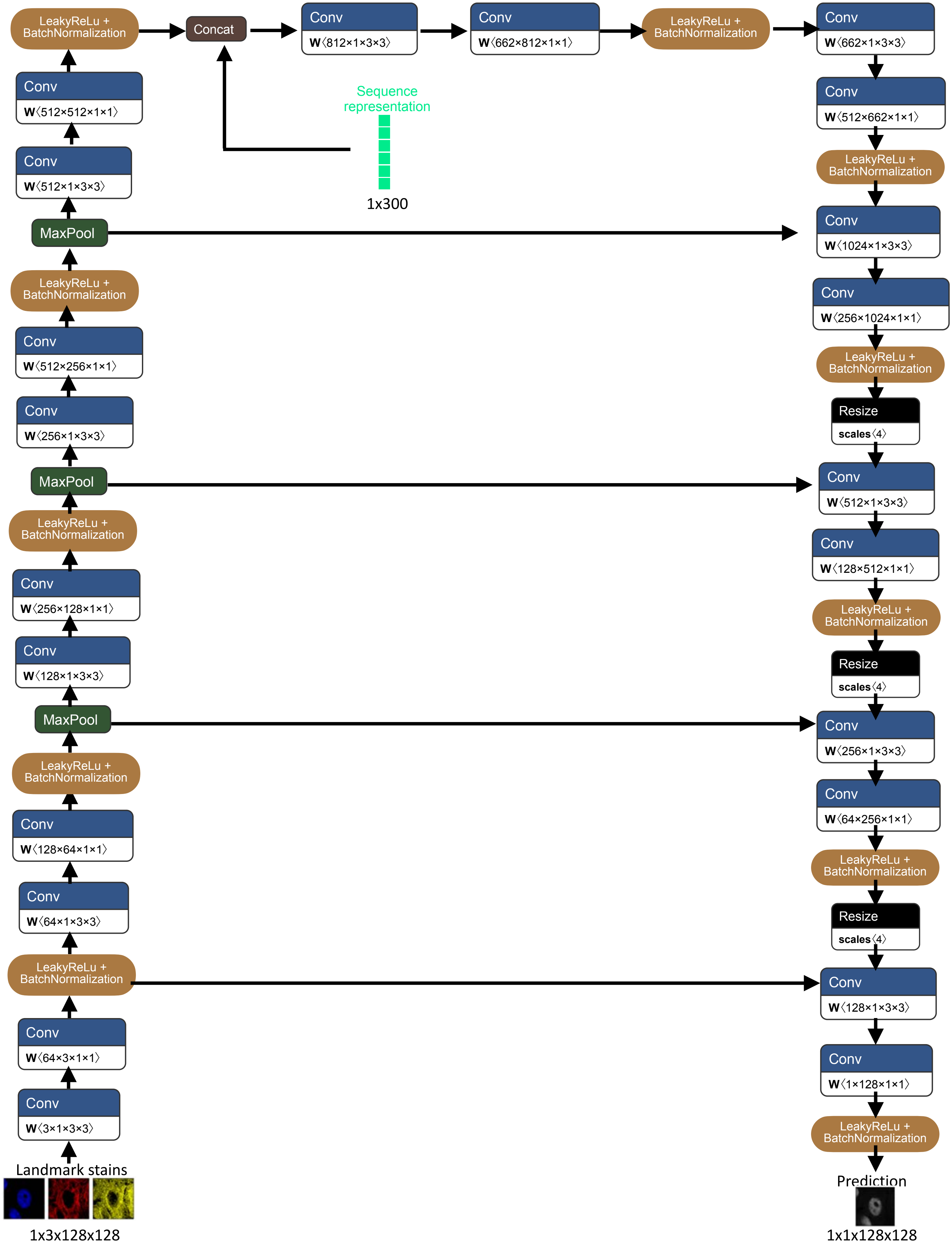


**Supplementary Figure 1**

**Supplementary Figure 1. Model architecture of image inpainting that learns an image representation from the cellular landmark stains and combines the protein sequence representation to predict the target protein image.**

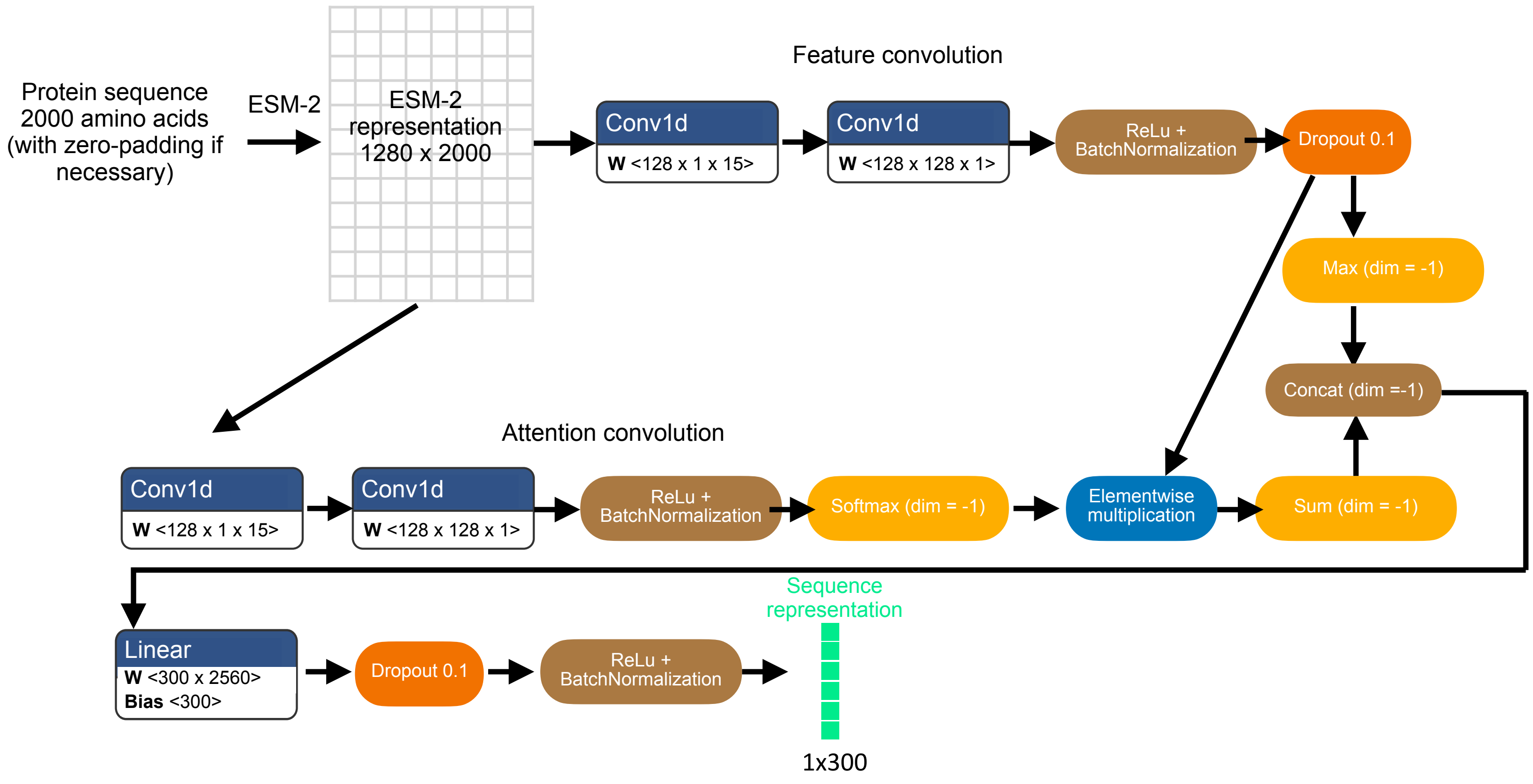


**Supplementary Figure 2**

**Supplementary Figure 2. Model architecture for learning protein sequence representation using the pretrained ESM-2 model[1] and a light attention layer[2].**

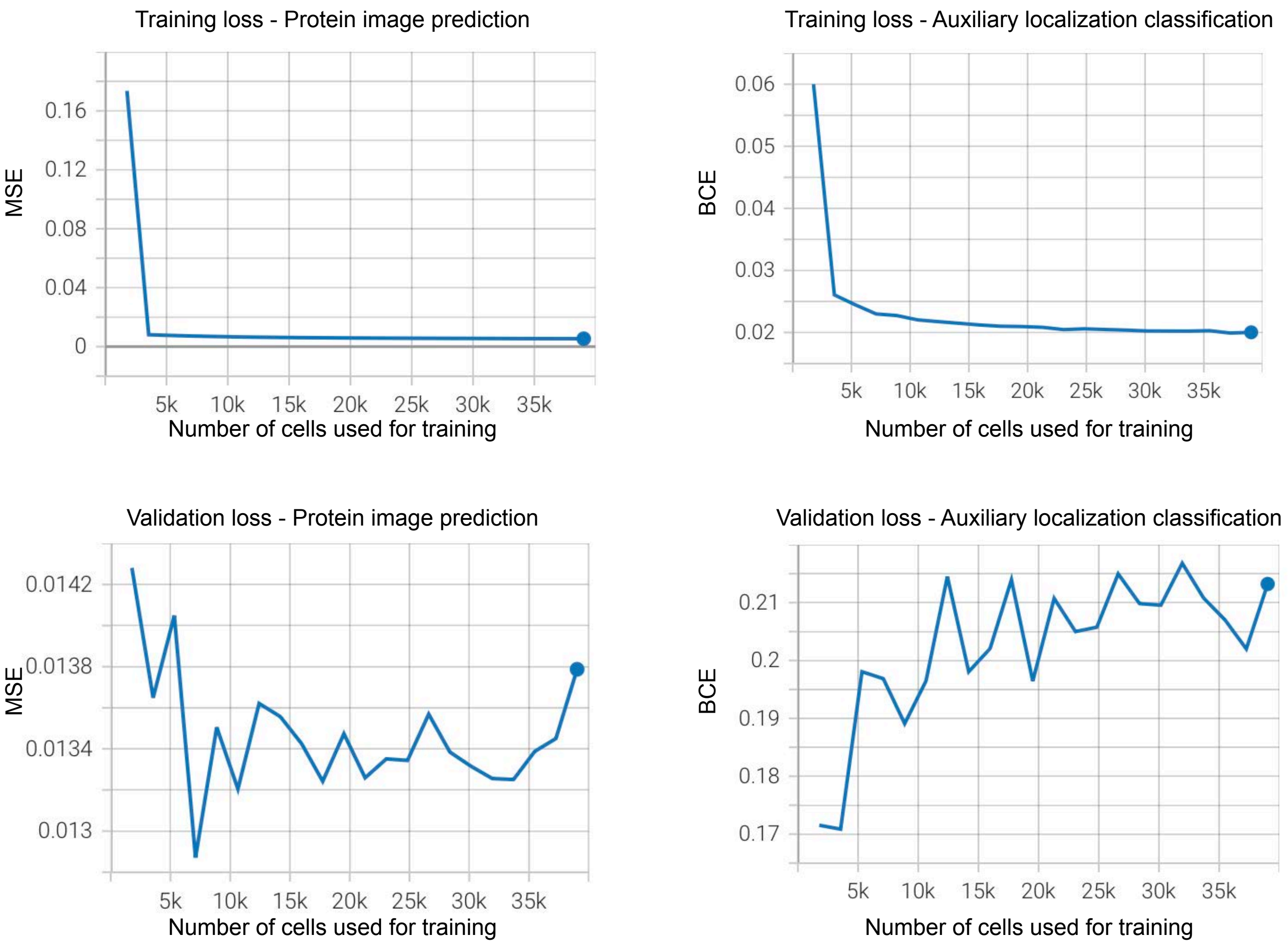


**Supplementary Figure 3**

**Supplementary Figure 3. Training and validation losses using the training dataset and the evaluation set of Holdout 1 (Figure 2a).**

**a** **Fluorescent spectrums overlap between the four channels.**

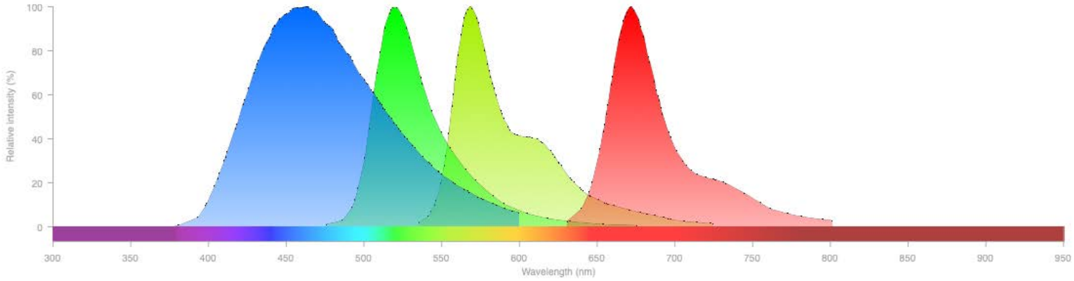


**b** **Intensity thresholding is necessary for model training.**

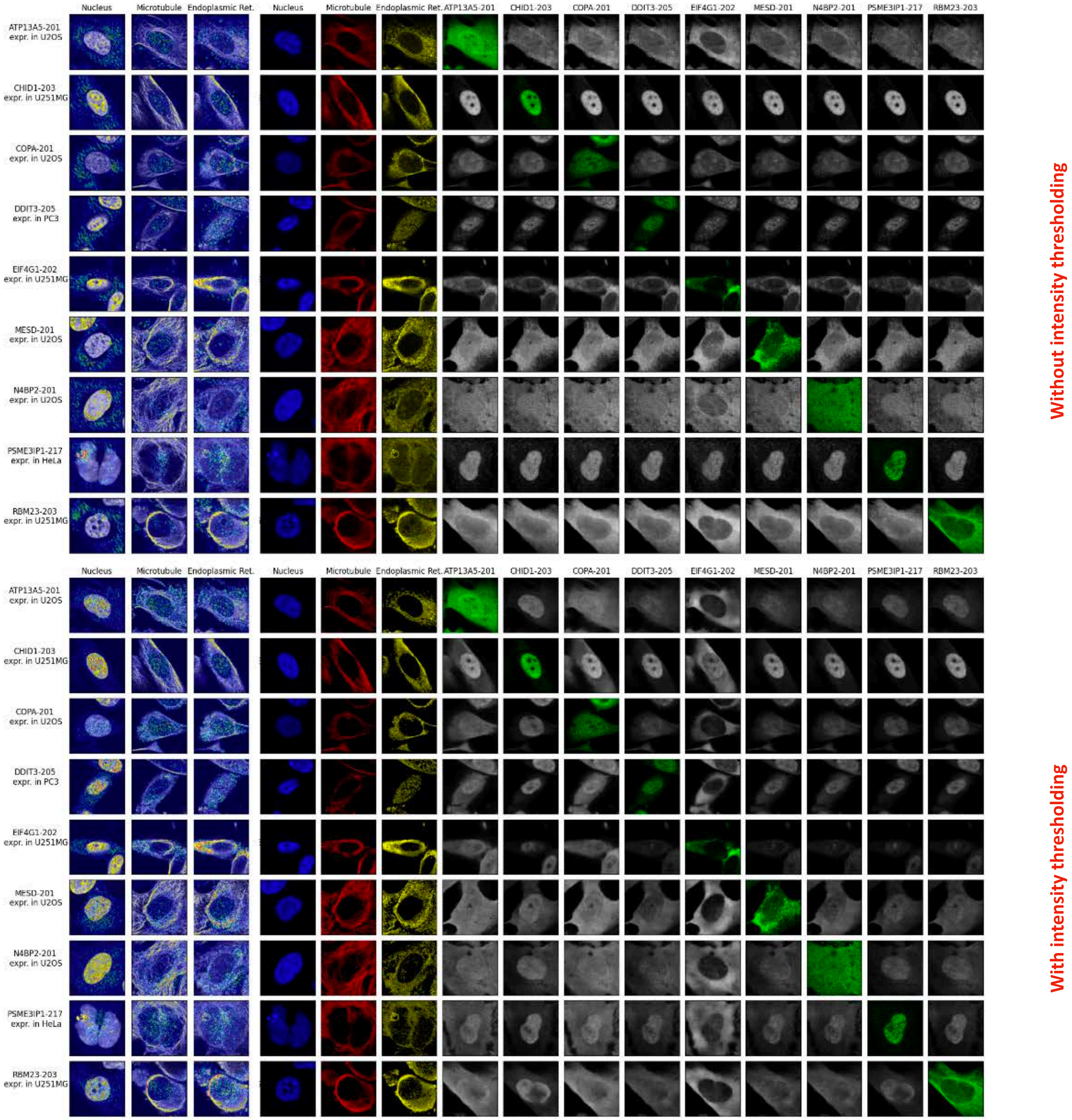


**Supplementary Figure 4**

**Supplementary Figure 4. Proper removal and/or assessment of spectral bleed-through is important for model training.**

(a) The emission spectrum of the four stains used in HPA are plotted (Alexa Fluor 555, 488, 647, and DAPI).

(b) We plotted the prediction of the proteins selected for experimental validation (Figure 5) in 9 cells in both the training set and Holdout 1 (Figure 2a). All predicted images are colored in grayscale and the green images on the diagonal are the real images from the HPA. The first three columns show the importance of each input pixel to the prediction of the protein images (green) on the diagonal, calculated using guided backpropagation[3]. The bottom panel shows the result of our model with the default setting, which has intensity thresholding at 0.19 (Methods), while the top panel shows the result of our model without any intensity thresholding. The model trained without thresholding (top) outputs images similar to the protein that was stained in HPA for the particular cell, regardless of the input protein sequence, whereas our default setting with intensity thresholding leads to different outputs for different proteins. Additionally, guided backpropagation[3] shows that the model without intensity thresholding pays attention to bleed-through signals outside of the staining target. For example, the last row of the top panel shows that the model is mostly using the signal outside of the nucleus in the DAPI channel to predict the RBM23 image.

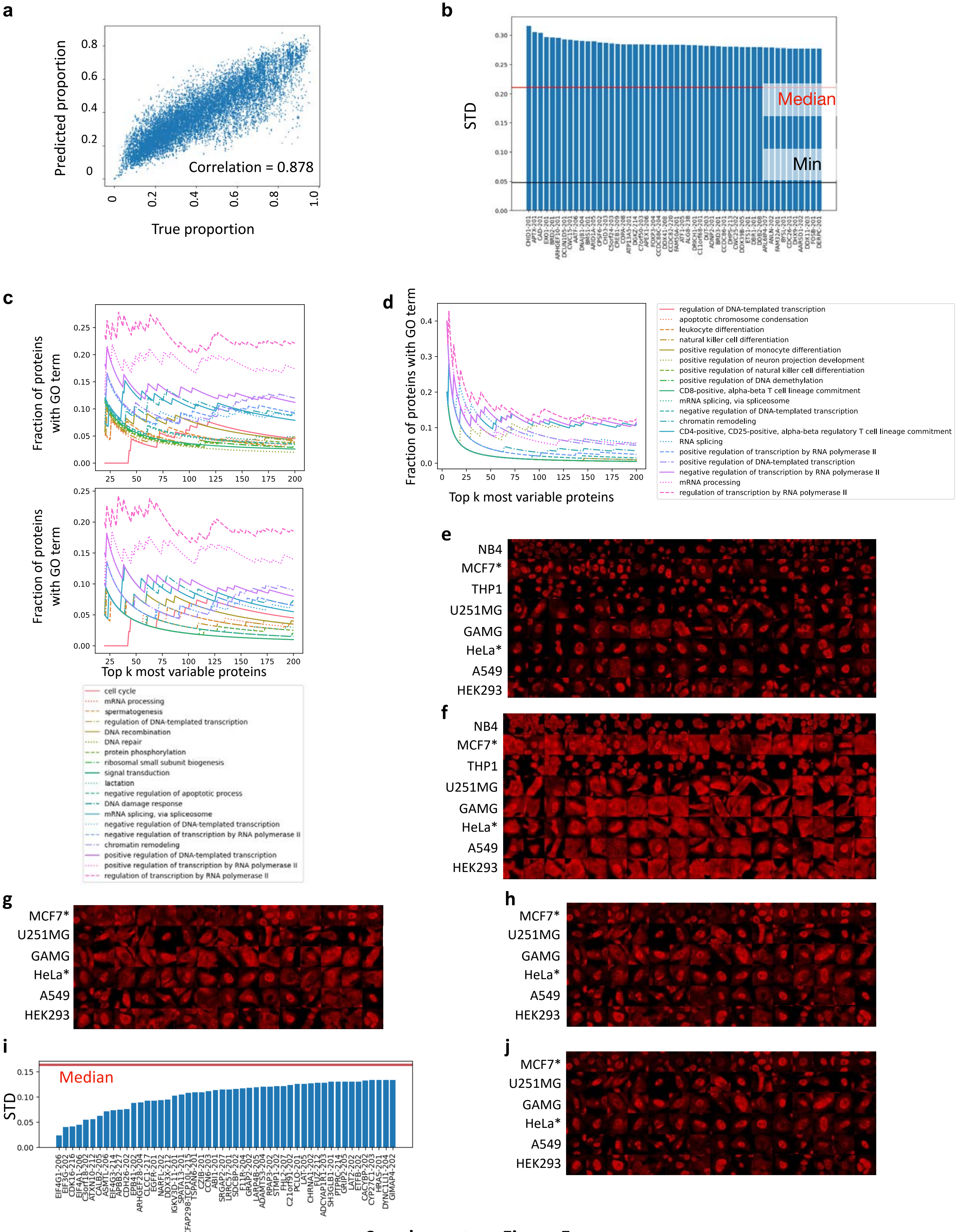


**Supplementary Figure 5**

**Supplementary Figure 5. Our model learns cellular contexts from landmark stains to accurately predict differences in protein localization across cell lines.**

(a) All held-out cells in the test set of Holdout 1 are plotted comparing the intra-nuclear proportion computed using the predicted protein images to the proportion computed using the real images from HPA. The Pearson correlation coefficient between the predicted intra-nuclear proportion and the experimentally measured proportion is 0.878. All cell lines and proteins used for plotting are not used in training the model and include proteins from unseen protein families that are not used in training.

(b) We computed the variability of the intra-nuclear proportion of a particular protein across cell lines. Standard deviation (STD) of the mean intra-nuclear proportion across 8 cell lines used in Figure 3 is computed for all proteins in the training set and Holdout 1 (Figure 2a). The proteins are ranked by their STD across the 8 cell lines. The red line indicates the median STD across all proteins. The black line indicates the minimum STD across all proteins.

(c) We plotted the gene ontology (GO) terms of the top k most variable proteins in Holdout 1 for k ranging from 20 to 200. For each value of k, we computed the fraction of genes annotated with each biological process term in GO. The most frequently occurring terms are indicated in the legend, from the least frequent to the most frequent. In order to visually separate the GO terms in the top panel, each term, from the least frequent (ranking = 0) to the most frequent (ranking = 18), is added with 0.005 * ranking of the term. Bottom panel shows the plot without the additional values added.

(d) Same plot as Figure 3c, without the additional 0.005 * ranking added for each GO term.

(e) For the 8 cell lines used in (b), we plotted the predicted images of one of the most variable proteins, CHID1-201 (in the training set) *: Cell lines are held-out from training the model. All cell crops are 50μm x 50μm.

(f) For the 8 cell lines used in (b), we plotted the predicted images of one of the most variable proteins, COPA-208 (in the training set) *: Cell lines are held-out from training the model. All cell crops are 50μm x 50μm.

(g) For 6 out of the 8 cell lines used in (b) that are contained in Holdout 2, we plotted the predicted images of one of the most variable proteins, MESD-201 (in Holdout 2) *: Cell lines are held-out from training the model. All cell crops are 50μm x 50μm.

(h) For 6 out of the 8 cell lines used in (b) that are contained in Holdout 2, we plotted the predicted images of one of the most variable proteins, RBM23-203 (in Holdout 2) *: Cell lines are held-out from training the model. All cell crops are 50μm x 50μm.

(i) Least variable proteins across the 6 selected cell lines in Holdout 2 are plotted (Methods).

(j) For 6 out of the 8 cell lines used in (b) that are contained in Holdout 2, we plotted the predicted images of one of the least variable proteins that are mainly inside the nucleus, PSME3IP1-217(in Holdout 2) *: Cell lines are held-out from training the model. All cell crops are 50μm x 50μm.

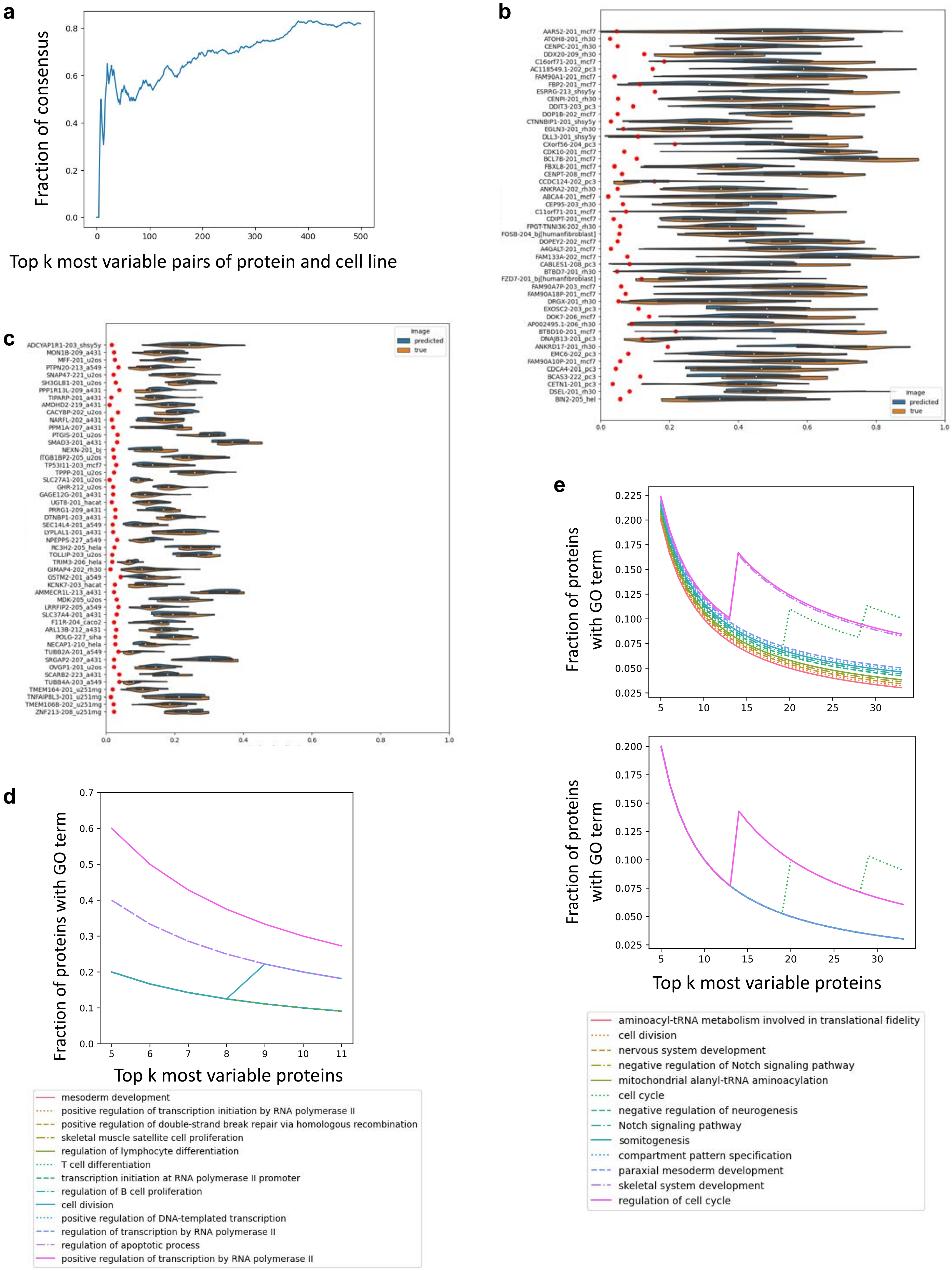


**Supplementary Figure 6**

**Supplementary Figure 6. Our model can predict variability in protein localization across single cells.**

(a) Protein-cell line pairs in Holdout 1 are ranked by the variance of intra-nuclear proportion. We compared this ranking computed from predicted protein images using held-out proteins and cell lines to the ranking computed from real protein images in the HPA. The fraction of protein-cell line pairs that overlap between the two rankings is plotted for the top k most variable pairs for k ranging from 1 to 500. All protein-cell line pairs used for plotting are not used in training the model and include proteins from unseen protein families that are not used in training.
(b) For the top 50 most variable pairs of protein-cell line pairs in Holdout 1 that overlap between the predicted and real HPA images, we plotted the distribution of intra-nuclear proportion for both the predicted and the true HPA images. Red dots indicate the average MSE loss of the predicted images compared to the true HPA images, demonstrating that the variability in intra-nuclear proportion is not a result of prediction errors.
(c) For the 50 least variable protein-cell line pairs in Holdout 2 that overlap between the predicted and real HPA images, we plotted the distribution of intra-nuclear proportion for both the predicted and the true HPA images. Red dots indicate the average MSE loss of the predicted images compared to the true HPA images.
(d) Same plot as Figure 4d, without the additional 0.005 * ranking added for each GO term.
(e) We plotted the gene ontology (GO) terms of the top k most variable proteins in Holdout 1. For each value of k, we computed the fraction of genes annotated with each biological process term in GO. The most frequently occurring terms are indicated in the legend, from the least frequent to the most frequent. In order to visually separate the GO terms in the top panel, each term, from the least frequent (ranking = 0) to the most frequent (ranking = 12), is added with 0.002 * ranking of the term. Bottom panel shows the plot without the additional values added.

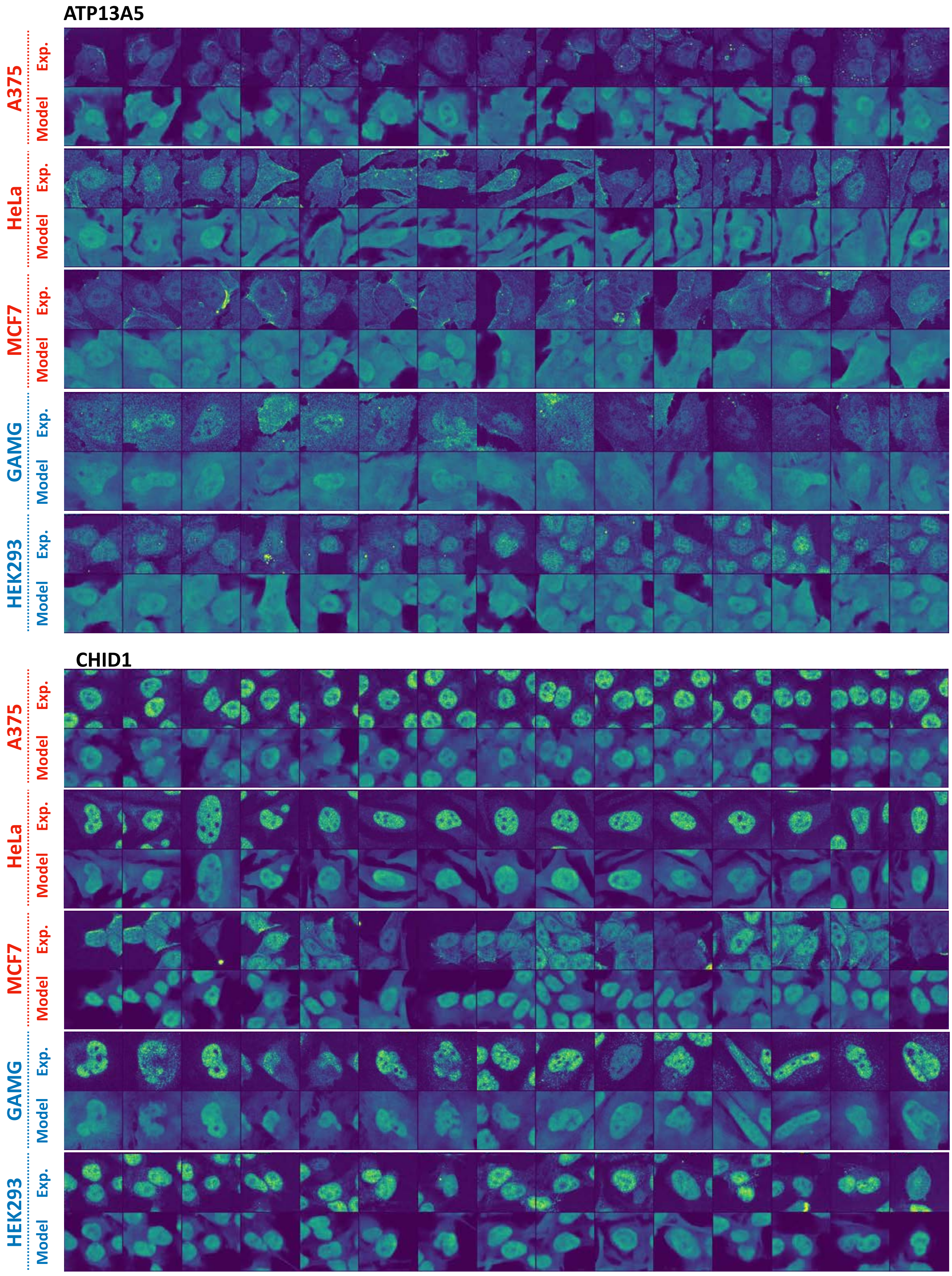


**Supplementary Figure 7**

**Supplementary Figure 7. Experimental validation of our model on 5 cell lines stained with ATP13A5 and CHID1.** 3 out of the 5 cell lines are not used in training the model (red). ATP13A5 and CHID1 were selected as the most variable proteins across cell lines (Methods). All cell crops are 50µm x 50µm.

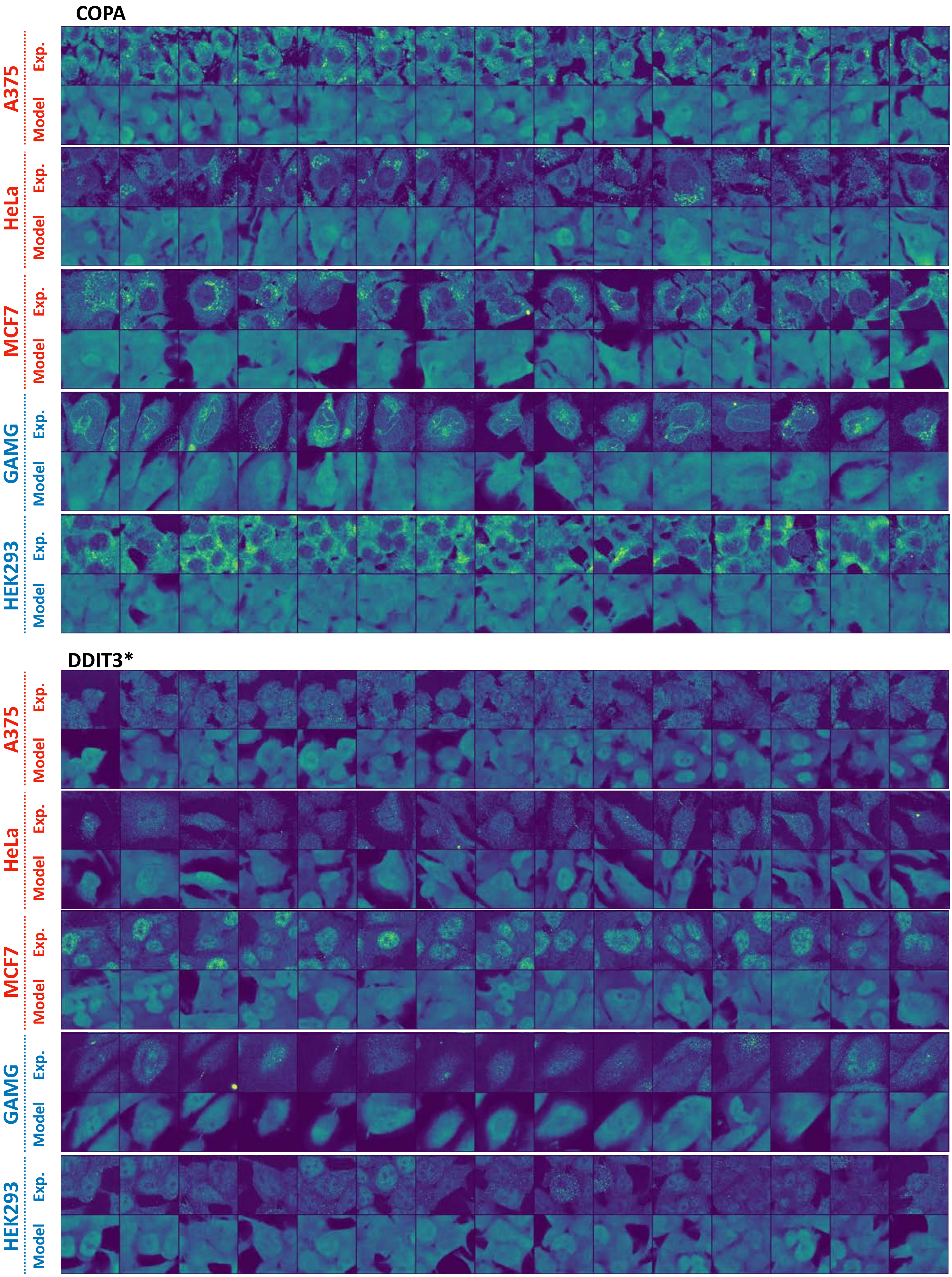


**Supplementary Figure 8**

**Supplementary Figure 8. Experimental validation of our model on 5 cell lines stained with COPA and DDIT3.** 3 out of the 5 cell lines are not used in training the model (red). *: DDIT3 is in the test set of Holdout 1. COPA was selected as one of the most variable proteins across cell lines (Methods). DDIT3 was selected as one of the most variable proteins across single cells of the same cell line (Methods). All cell crops are 50μm x 50μm.

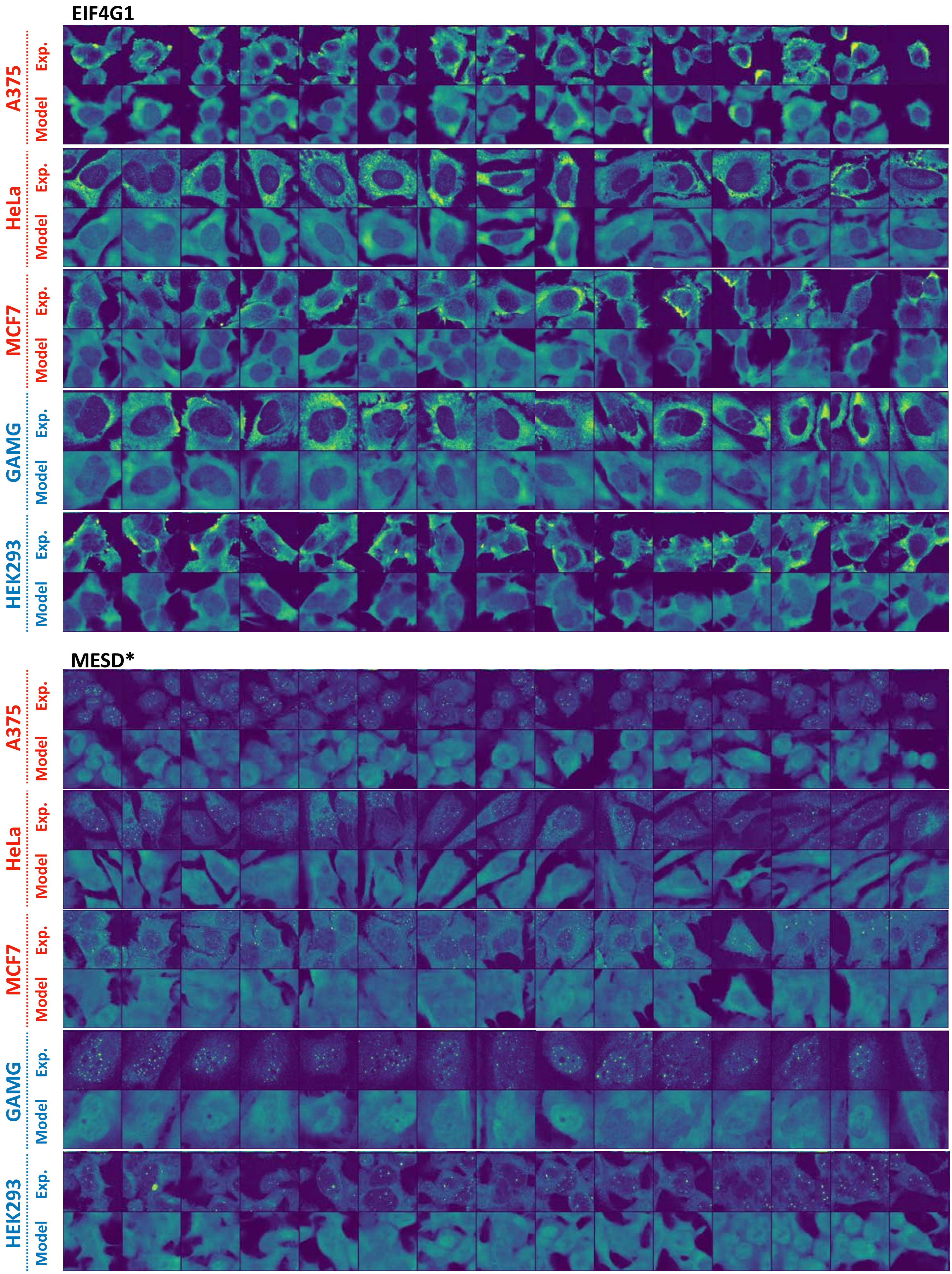


Supplementary Figure 9

**Supplementary Figure 9. Experimental validation of our model on 5 cell lines stained with EIF4G1 and MESD.** 3 out of the 5 cell lines are not used in training the model (red). *: MESD is in Holdout 2. MESD was selected as one of the most variable proteins across cell lines (Methods). EIF4G1 was selected as one of the least variable proteins across cell lines and is predicted to be localized mainly outside of the nucleus. All cell crops are 50μm x 50μm.

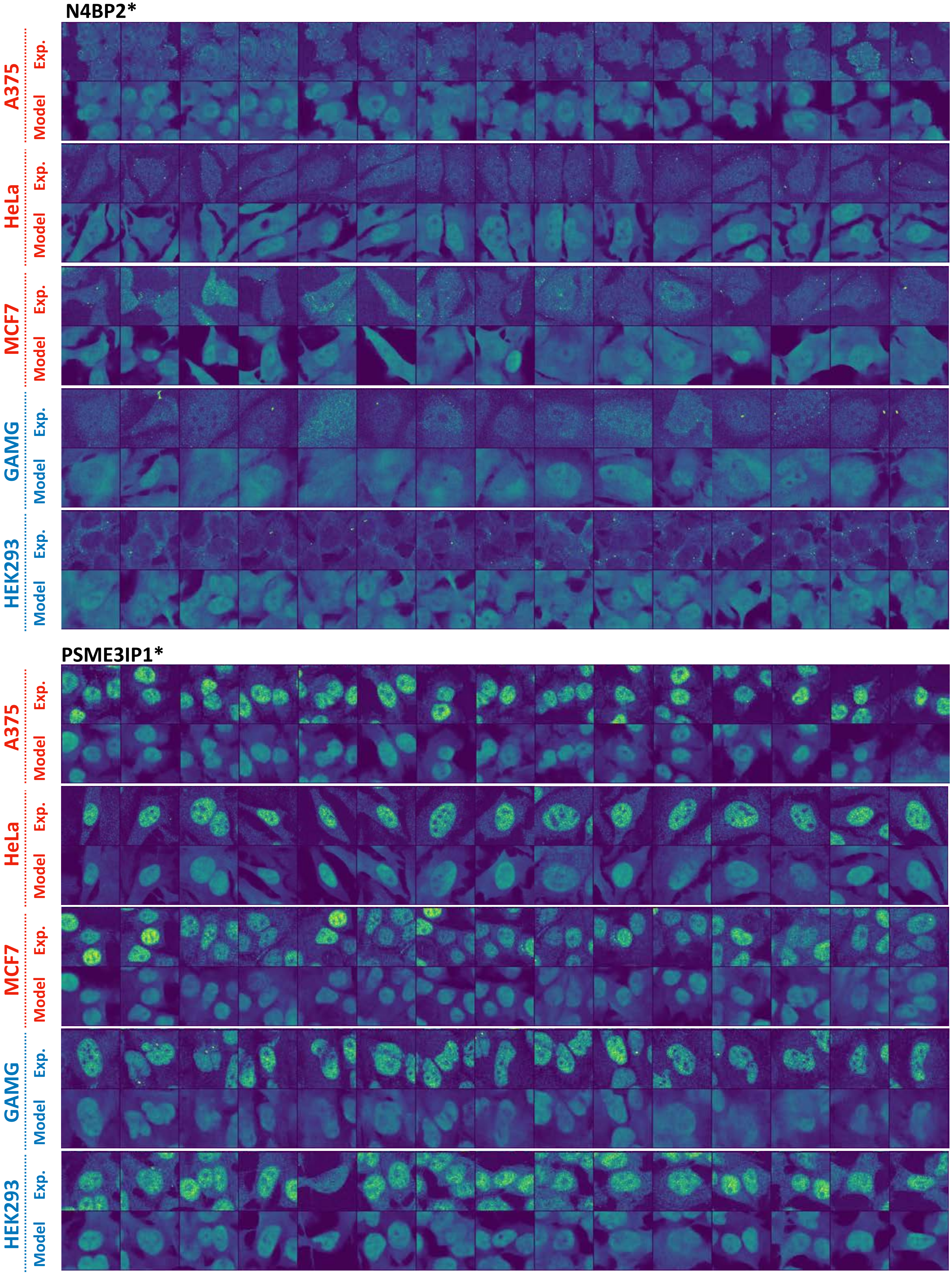


**Supplementary Figure 10**

**Supplementary Figure 10. Experimental validation of our model on 5 cell lines stained with N4BP2 and PSME3IP1.** 3 out of the 5 cell lines are not used in training the model (red). *: Both proteins are in Holdout 2. N4BP2 was selected as one of the most variable proteins across single cells of the same cell line. PSME3IP1 was selected as one of the least variable proteins across cell lines and is predicted to be localized mainly inside the nucleus. All cell crops are 50μm x 50μm.

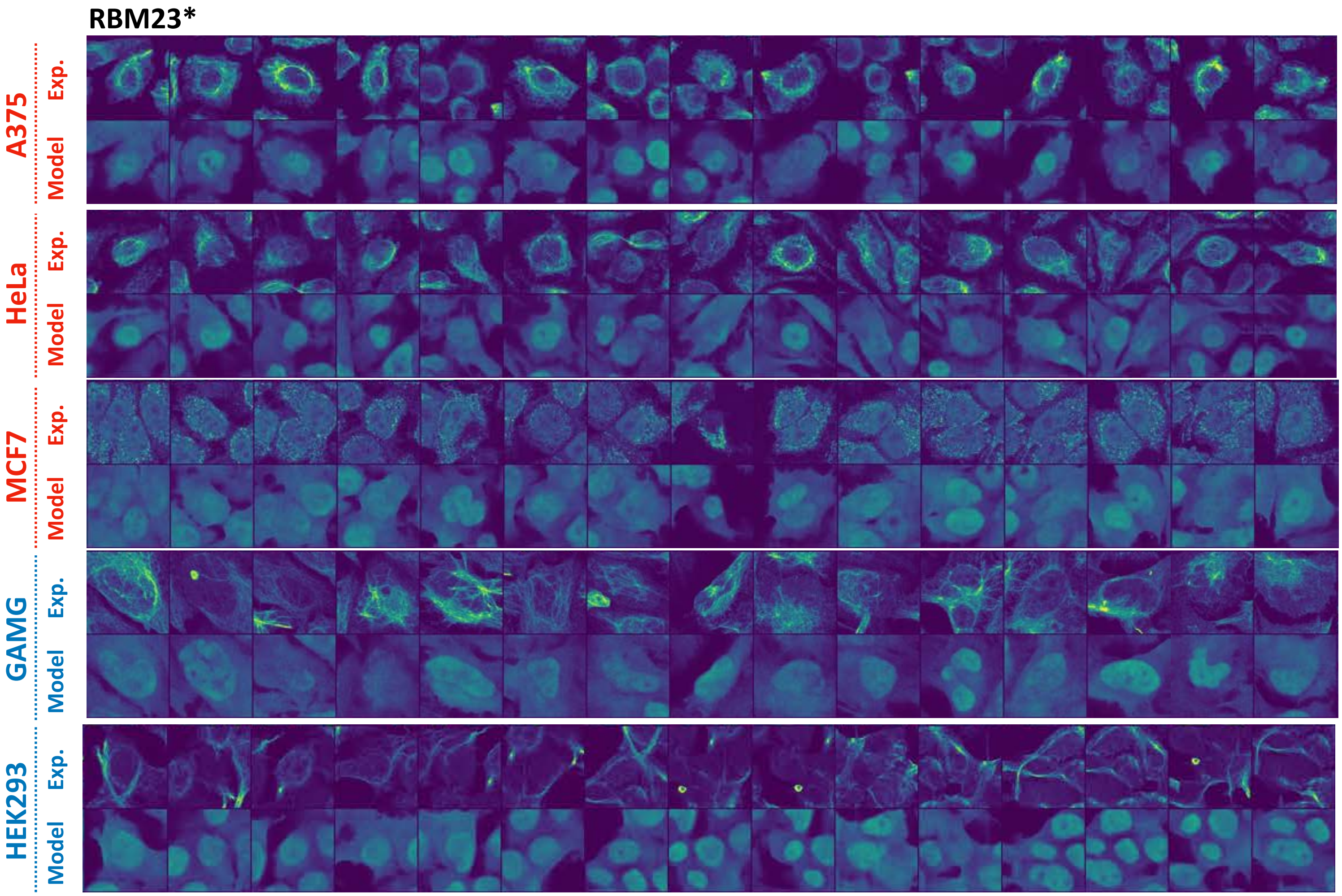


**Supplementary Figure 11**

**Supplementary Figure 11. Experimental validation of our model on 5 cell lines stained with RBM23.** 3 out of the 5 cell lines are not used in training the model (red). *: RBM23 is in Holdout 2. RBM23 was selected as one of the most variable proteins across cell lines. All cell crops are 50μm x 50μm.

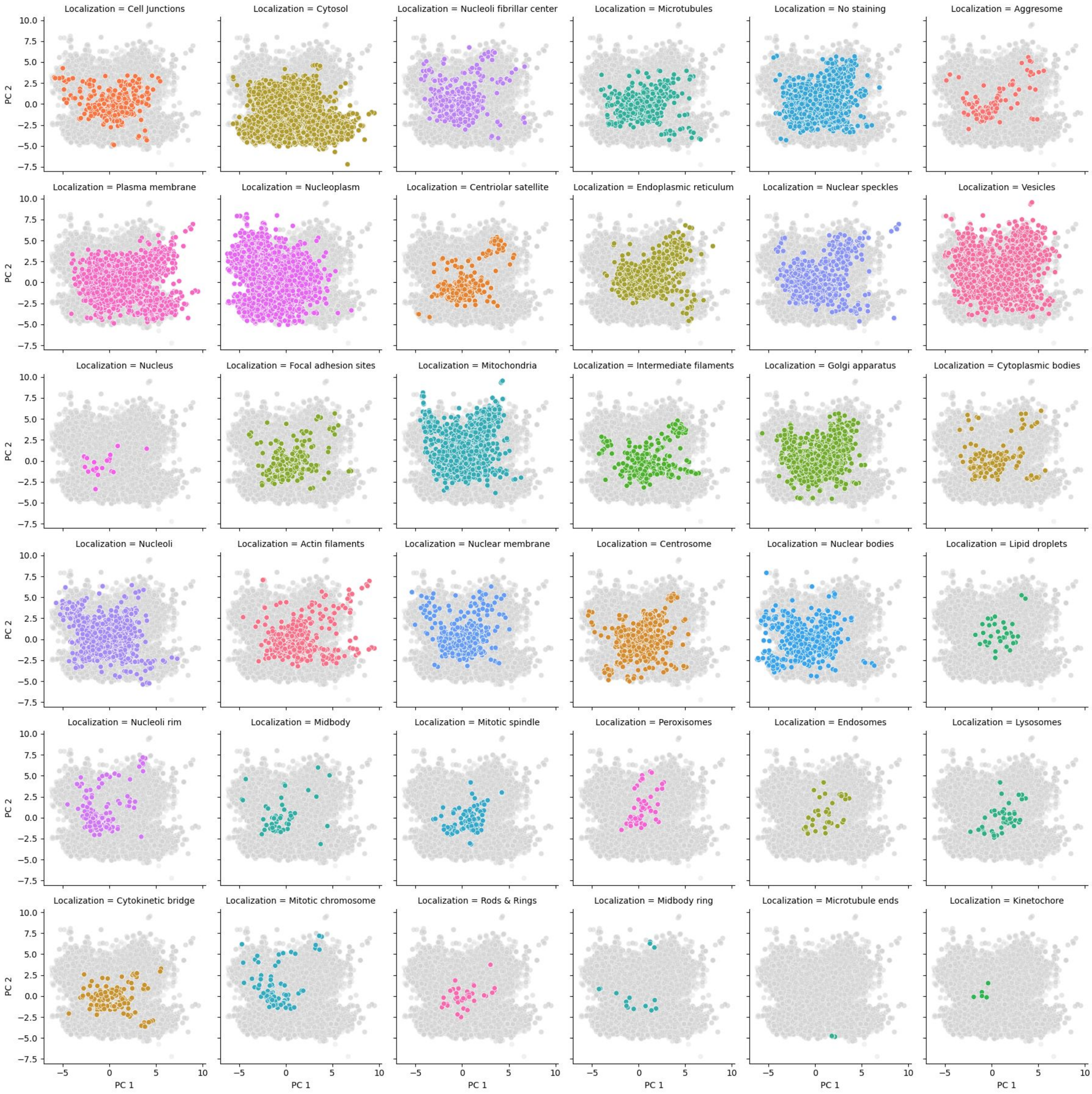


**Supplementary Figure 12**

**Supplementary Figure 12. PCA of protein sequence representation of 40,622 proteoforms corresponding to 12,614 genes.**

We used the same PCA embeddings of protein sequences as shown in Figure 6a. Each panel shows the sequence representation of proteoforms annotated with a particular localization, while all other proteoforms are colored in gray.

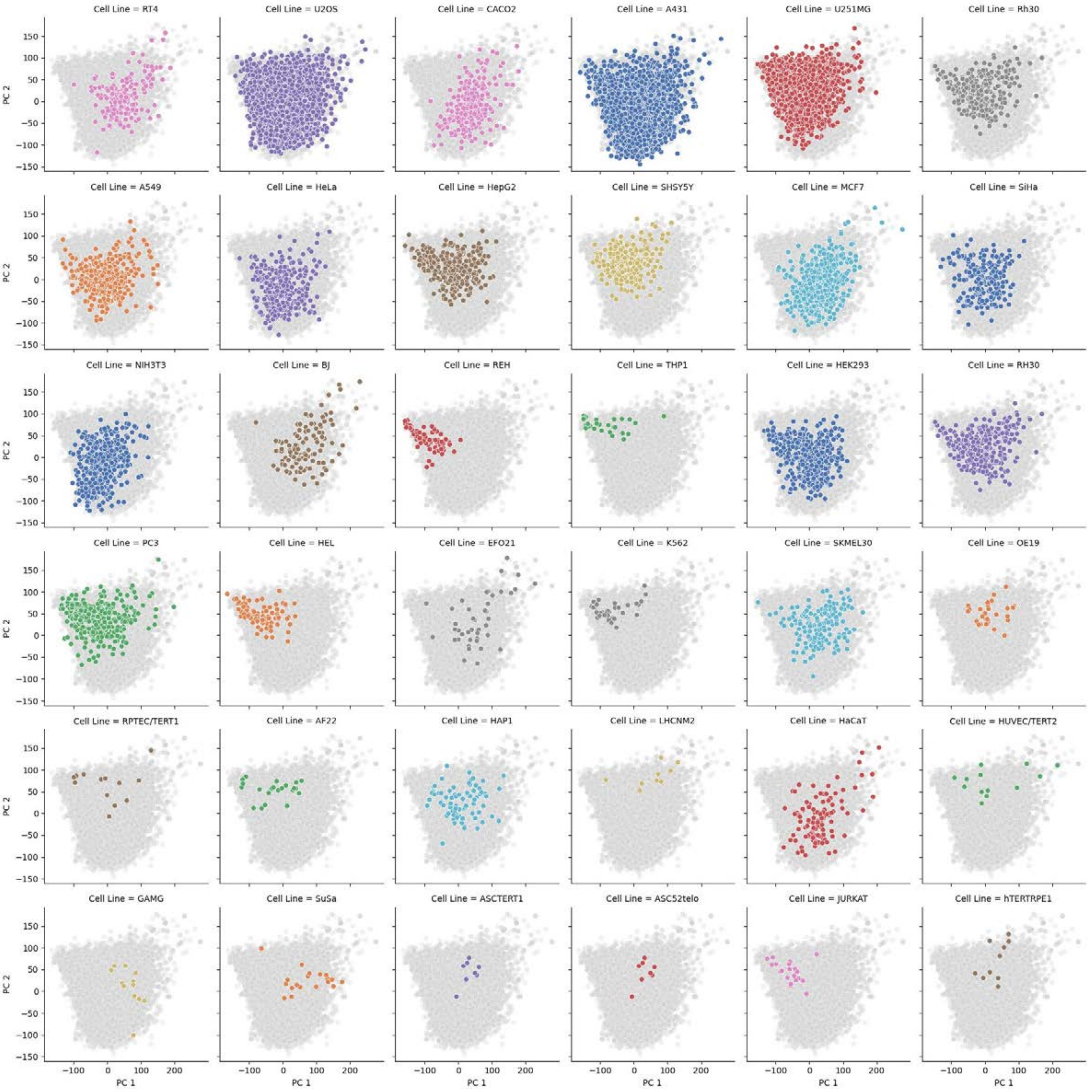


**Supplementary Figure 13**

**Supplementary Figure 13. PCA of 36 cell lines.**

We computed PCA of the image representations (Figure 1b) of the landmark stains from 36 cell lines. Each panel shows the image representations of cells in a particular cell line, while all other cell lines are colored in gray.

| | Name | Clone | Vendor | Catalog # | Host | Stock | Final concentration |
|---|---|---|---|---|---|---|---|
| **1st Ab** | a-Tubulin | DM1A | Abcam | ab7291 | Mouse | 1.0 mg/mL | 1.0 µg/mL |
| **1st Ab** | calreticulin | poly | Abcam | ab2908 | Chicken | 1.0 mg/mL | 1.25 µg/mL |
| **1st Ab** | EIF4G1 | HPA028487 | Atlas Antibodies | | Rabbit | 0.1 mg/mL | 2.0 µg/mL |
| **1st Ab** | DDIT3 | HPA058416 | Atlas Antibodies | | Rabbit | 0.3 mg/mL | 2.0 µg/mL |
| **1st Ab** | N4BP2 | HPA042607 | Atlas Antibodies | | Rabbit | 0.2 mg/mL | 2.0 µg/mL |
| **1st Ab** | CHID1 | HPA039374 | Atlas Antibodies | | Rabbit | 0.05 mg/mL | 2.0 µg/mL |
| **1st Ab** | COPA | HPA028024 | Atlas Antibodies | | Rabbit | 0.1 mg/mL | 2.0 µg/mL |
| **1st Ab** | ATP13A5 | HPA031772 | Atlas Antibodies | | Rabbit | 0.3 mg/mL | 2.0 µg/mL |
| **1st Ab** | MESD (MESDC2) | HPA039414 | Atlas Antibodies | | Rabbit | 0.1 mg/mL | 2.0 µg/mL |
| **1st Ab** | RBM23 | HPA004144 | Atlas Antibodies | | Rabbit | 0.1 mg/mL | 2.0 µg/mL |
| **1st Ab** | PSME3IP1 (FAM192A) | HPA054382 | Atlas Antibodies | | Rabbit | 0.1 mg/mL | 2.0 µg/mL |
| **2nd Ab** | Anti-rabbit Alexa488 | Poly | Thermo | A11034 | Goat | 2.0 mg/mL | 2.5 µg/mL |
| **2nd Ab** | Anti-mouse Alexa555 | Poly | Thermo | A21424 | Goat | 2.0 mg/mL | 2.5 µg/mL |
| **2nd Ab** | Anti-chicken Alexa647 | Poly | Thermo | A21449 | Goat | 2.0 mg/mL | 2.5 µg/mL |
| **DAPI** | DAPI | N/A | Sigma | F0895 | N/A | 0.4 µg/mL | 0.2 µg/mL |

**Supplementary Table 1**

**Supplementary Table 1. Antibodies used for experimental validation.**
